# Soil microbial community response to corrinoids is shaped by a natural reservoir of vitamin B_12_

**DOI:** 10.1101/2024.02.12.580003

**Authors:** Zachary F. Hallberg, Alexa M. Nicolas, Zoila I. Alvarez-Aponte, Kenny C. Mok, Ella T. Sieradzki, Jennifer Pett-Ridge, Jillian F. Banfield, Hans K. Carlson, Mary K. Firestone, Michiko E. Taga

**Affiliations:** Department of Plant & Microbial Biology, University of California, Berkeley, Berkeley, CA, 94720 USA; Environmental Science, Policy and Management, University of California, Berkeley, Berkeley, CA, 94720 USA; Lawrence Livermore National Laboratory, Livermore, CA, 94550 USA; Innovative Genomics Institute, Berkeley, CA, 94720 USA; Lawrence Berkeley National Laboratory, Berkeley, CA, 94720 USA; Earth and Planetary Science, University of California, Berkeley, Berkeley, CA, 94720 USA

**Keywords:** corrinoids, cobalamin, vitamin B12, microbiome, soil, enrichments, bacteria, archaea, cofactor

## Abstract

Soil microbial communities perform critical ecosystem services through the collective metabolic activities of numerous individual organisms. Most microbes use corrinoids, a structurally diverse family of cofactors related to vitamin B_12_. Corrinoid structure influences the growth of individual microbes, yet how these growth responses scale to the community level remains unknown. Analysis of metagenome-assembled genomes suggests corrinoids are supplied to the community by members of the archaeal and bacterial phyla *Thermoproteota*, *Actinobacteria*, and *Proteobacteria*. Corrinoids were found largely adhered to the soil matrix in a grassland soil, at levels exceeding those required by cultured bacteria. Enrichment cultures and soil microcosms seeded with different corrinoids showed distinct shifts in bacterial community composition, supporting the hypothesis that corrinoid structure can shape communities. Environmental context influenced both community and taxon-specific responses to specific corrinoids. These results implicate corrinoids as key determinants of soil microbiome structure and suggest that environmental micronutrient reservoirs promote community stability.

## Introduction

With thousands of bacterial species and billions of cells per gram, soils are believed to harbor the most diverse microbial communities, and are perhaps the final frontier of environmental microbiology (ref. 1). The complexity of soil presents challenges for the characterization of microbial functions *in situ*, yet recent-omics and biochemical studies have revealed succession patterns of microbial populations in soils across growing seasons (ref. 2) and how soil microbial communities respond to environmental factors ranging from temperature to nutrient availability (ref. 3, 4). These studies demonstrate that soil microbial communities are dynamic in response to environmental changes, and that understanding how these communities respond to specific perturbations is essential to promoting their health and function.

Microbial communities can be modulated by addition of limiting nutrients, evidenced by soil microbiome responses to the addition of nutrients such as nitrogen and phosphorus (ref. 3, 4, 5). These nutrients become assimilated into biomass and are chemically transformed as they cycle through the community. Comparatively little is known about microbiome responses to those nutrients that are not incorporated into biomass, such as enzyme cofactors (ref. 6, 7, 8). It has long been established that many soil bacteria depend upon the biosynthetic capacity of other organisms to provide key cofactors necessary for their growth, such as B vitamins (ref. 9, 10, 11). However, the impact of interactions involving enzyme cofactors on soil microbiome composition and function remains unknown.

One exemplary class of shared nutrients is corrinoids, the vitamin B_12_ (cobalamin) family of cobalt-containing modified tetrapyrroles used as cofactors by the majority of organisms (Fig. 1A) (ref. 12). Corrinoid-dependent metabolism is widespread across bacteria: 86% of publicly available bacterial genomes contain one or more corrinoid-dependent enzymes involved in nucleotide metabolism (e.g. ribonucleotide reductase [RNR]), amino acid metabolism (e.g methionine synthase), carbon catabolism (e.g. methylmalonyl-CoA mutase [MCM] and methyltransferases involved in one-carbon metabolism), and other processes (ref. 12, 13, 14, 15). However, only 37% of these genomes encode the *de novo* corrinoid biosynthesis pathway, suggesting that corrinoids are widely shared in communities (Fig. 1B) (ref. 13, 14, 16). This genomic evidence is corroborated by experimental studies showing that corrinoids are key shared nutrients in ocean and animal gut environments (ref. 14, 17) and in cultured microbes (ref. 10, 14, 18).

**Figure 1.**
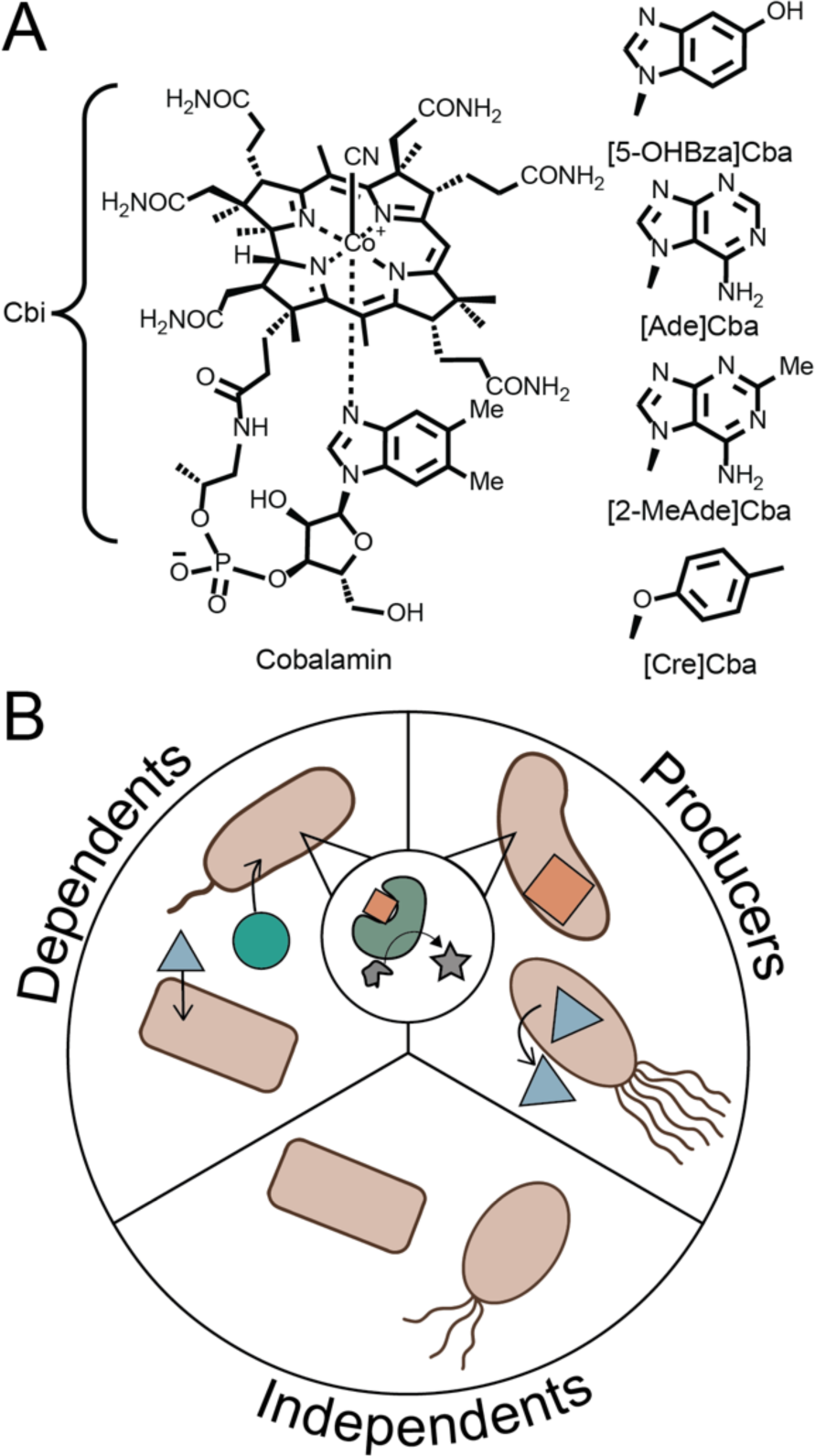
Corrinoids and corrinoid ecological roles. A. Structure of cyanocobalamin (vitamin B12) and related corrinoids used in this study. Cobinamide (Cbi) is a truncated precursor. Other corrinoids used in this study differ from cobalamin in the structure of the lower ligand. B. Ecological roles of microbes with respect to corrinoids. Microbes may biosynthesize a corrinoid (“Producers”), use but not produce a corrinoid (“Dependents”), or possess no corrinoid biosynthetic enzymes or requirements (“Independents”).

Mounting evidence suggests that corrinoids are also critical nutrients in soil. Early research demonstrated that cobalamin could substitute for extracts from soil in stimulating growth and cultivation of soil bacteria (ref. 10, 19). Indeed, cobalamin has been detected at levels ranging from 1 to 50 ng per gram of soil in 11 soils in Poznań, Poland, using a microbial growth-based assay (ref. 20). The presence of cobalamin in soil was substantiated by LC-MS analysis in over 40 diverse soil types across Canada and Scotland (ref. 21). This study further found through sequence analysis that genera containing corrinoid-dependent enzymes account for at least 70% of the total detected soil community across 155 metagenomes, whereas less than 10% of genera encode genes for *de novo* corrinoid biosynthesis (ref. 21). More recently a survey of marine and soil metagenome-assembled genomes (MAGs) showed that predicted corrinoid-using microorganisms dominate globally distributed soils (ref. 22). The past 70 years of findings suggest that corrinoids are widely used and shared metabolites in soil microbial communities.

Unlike other enzyme cofactors, corrinoid cofactors are structurally diverse. Cobalamin has historically received significant attention in biomedical research as it is considered the only corrinoid used by humans (ref. 23). However, nearly 20 different corrinoids with variations in the lower ligand structure have been identified in cultured microbes and microbial communities (Fig. 1A) (ref. 24, 25, 26, 27, 28). The influence of corrinoid lower ligand structure on enzyme function, microbial growth, and community structure is rarely studied systematically because corrinoid cofactors other than cobalamin are not commercially available. Experimental studies using corrinoids extracted and purified from microbial cultures have shown that the structure of the lower ligand impacts corrinoid uptake, enzyme binding, and microbial growth in pure culture (ref. 29, 30, 31, 32, 33). The structural specificity of corrinoid-based interactions also extends to coculture-based studies, where corrinoid type can either promote or prevent the growth of one or both members of the culture (ref. 29, 32, 34). Because corrinoids are required by many organisms, structurally diverse, and functionally specific, they are ideal models for shared nutrients (ref. 35).

Despite the growing appreciation that corrinoids with chemically distinct lower ligands differentially impact axenic microbial functions and growth, it remains unknown if and how these direct effects on individual organisms scale to highly complex communities such as those of soil. Here, we use corrinoids as a model nutrient to investigate the importance and influence of corrinoids in a California grassland soil. We find that corrinoid-dependent metabolism is widespread, whereas predicted capacity for *de novo* biosynthesis is restricted, suggesting that corrinoid sharing interactions are prevalent. We find that this soil contains an abundance of cobalamin, which may explain why the resident microbial community is resilient to additional cobalamin unless the community is removed from the soil through enrichment culturing. In contrast, corrinoids undetected in this soil can profoundly, but transiently, shift the community both in soil and in enrichment communities. Our results demonstrate that shared, non-assimilated nutrients represent an underappreciated component of soil microbiome function, and provide a blueprint for further investigation of corrinoids or other shared nutrients in this system.

## Materials and Methods

### Establishing corrinoid use and dependence of a soil MAG set

503 dereplicated MAGs from soil sampled at the University of California operated Hopland Research and Extension Center in Hopland, CA (39.004056 N, 123.085861 w) in February, March (ref. 36), and August (ref. 37, 38) 2018 were analyzed for their corrinoid-related ecological roles. MAGs previously annotated using DRAM (ref. 37, 38, 39) were cross-listed with KEGG orthologues (KOs) (unless otherwise stated) for cobamide annotations outlined in (ref. 16). MAGs were inspected for their potential capacity to produce corrinoids based on the presence of one or more marker genes that are 99% correlated with *de novo* corrinoid biosynthesis in publicly available genomes (ref. 16), and which we have validated in two experimental studies (ref. 16, 18). These six marker genes correspond to three enzymatic steps in corrin ring biosynthesis that have anaerobic and aerobic orthologs: *cbiC* (K06042), *cbiF* (K05936), *cbiL* (K03394), *cobH* (K06042), *cobI* (K03394), *cobM* (K05936). We also used KOs to identify corrinoid-independent and -dependent genes in MAGs. The single exception was the KOfam K00525 for class II (corrinoid-dependent) RNR which, after manual inspection of annotations, we found incorrectly includes both the class II and class I (corrinoid-independent) RNRs. We therefore used the Pfam PF08471, which differentiated the class II RNR. We defined corrinoid ecological roles based on the number of unique corrinoid-dependent and corrinoid biosynthesis marker genes per genome. MAGs that contained at least one biosynthesis marker gene and at least one corrinoid-dependent gene were defined as “producers”. Those with one or more biosynthesis marker gene and no corrinoid-dependent gene were defined as “producer/non-users.” Both “dependent” and “independent” MAGs were those with no biosynthesis marker gene; those that contained at least one corrinoid-dependent gene were assigned to the dependent category, and those with none were categorized as independent. Further manual searches were performed for the *Nitrososphaera* MAG identified as a producer/non-user. This genome contained two annotated cobalt importers, corrinoid-independent RNRs, and the genes of the corrinoid-independent ethanolamine utilization pathway. We combed its ORFs for other cobalamin- or corrinoid-use annotations in addition to corrinoid binding domains (via Pfams PF02310 and PF02607) and could not find evidence of corrinoid use. All corrinoid annotations used in this analysis can be found in Supp. Table 1. Although high-affinity corrinoid uptake is presumably required for corrinoid acquisition by dependents (ref. 14), the limited annotations for transporters prevented us from making reliable predictions of corrinoid uptake genes (ref. 16).

### Soil Sampling

We collected soil samples from the University of California operated Hopland Research and Extension Center in Hopland, CA (39.004056 N, 123.085861 W) on April 22, 2021. Soil was sampled from the top 10 cm and sieved in the field to 2 mm. Samples used for laboratory microcosms were pre-wetted to 20% field moisture 1 week before beginning the experiment.

### Liquid-solid extraction of soil using organic solvents

Extraction solutions (50 mL) were prepared fresh from methanol (Fisher Scientific, Pittsburgh, PA, USA), acetonitrile (Fisher Scientific, Pittsburgh, PA, USA), chloroform (Fisher Scientific, Pittsburgh, PA, USA), ethyl acetate (VWR, Radnor, PA, USA), formic acid (Fisher Scientific, Pittsburgh, PA, USA), ammonium hydroxide (Fisher Scientific, Pittsburgh, PA, USA), and double-distilled H_2_O (ddH_2_O). Extraction solutions were composed of: Solution A – MeOH:H_2_O (4:1), Solution B – CHCl_3_:MeOH (2:1), Solution C – EtOAc, Solution D – Acidic MeOH:H_2_O (4:1), Solution E – Basic MeOH:H_2_O (4:1), Solution F – Acidic MeCN:MeOH:H_2_O (2:2:1), Solution G – Basic MeCN:MeOH:H_2_O (2:2:1), Solution H – MeCN:MeOH:H_2_O (2:2:1), Solution I – Acidic MeCN:H_2_O (4:1), Solution J – Basic MeCN:H_2_O (4:1), Solution K – MeCN:H_2_O (4:1). Acidic extraction solutions included 0.1 M formic acid, and basic solutions included 0.1 M ammonium hydroxide.

Each extraction was performed as follows: to a 15-mL polystyrene centrifuge tube (Falcon, Corning, Glendale, AZ), 2 g of soil were added. Extraction solution (5 mL) was added to the soil and vortexed for 1 minute at room temperature, followed by incubation at room temperature for 10 minutes. The extraction tube was centrifuged at 7,197 × rcf for 2.5 minutes, and the supernatant decanted into a 20 mL scintillation vial. This extraction was repeated 2 additional times, for a total of 3 extraction rounds per extraction. Extract supernatants were then filtered through a 0.2 µm nylon filter (Pall Corporation, Port Washington, NY), and evaporated to dryness with a rotary evaporator (Heidolph Collegiate, Wood Dale, IL).

Dried extract samples were resuspended in 500 µL ddH_2_O, transferred to a 1.7 mL microcentrifuge tube (VWR, Radnor, PA), heated to 95°C for 15 minutes, cooled to room temperature, and centrifuged for 15 min at 20,187 × rcf.

### E. coli corrinoid quantification bioassay

The corrinoid quantification bioassay and strains are described previously (ref. 40). *E. coli* strains MG1655 (wild type, WT) and isogenic *ΔmetE* and *ΔmetE ΔmetH* were streaked from 25% glycerol freezer stocks onto LB-agar (1.5%) plates to obtain single colonies. M9-glycerol (0.4%) minimal medium supplemented with 6.7 mM L-methionine (5 mL) in a culture tube was inoculated with cells from a single colony of the bioassay strain, and grown for 24 h at 37°C with shaking at 210 rpm. Cells were pelleted by centrifugation (4000 × rcf, 4 min) of 1 mL of saturated culture, the supernatant aspirated, and resuspended in 1×PBS (1 mL, composed of 10.1 mM Na_2_HPO_4_, 1.8 mM KH_2_PO_4_, 137 mM NaCl, 2.7 mM KCl, pH 7.4). Cells were washed in this way twice more, and then diluted 10-fold in 1×PBS prior to use.

For corrinoid measurements, 2 µL of washed cells were added to a mixture of 180 µL M9-glycerol (0.4%) medium and 20 µL control or experimental solution. Control solutions consisted of L-methionine (final concentration 1 mM), cyanocobalamin (CN-Cbl) or dicyanocobinamide (CN_2_-Cbi) (final control concentrations 1 nM, purchased from MilliporeSigma, Burlington, MA), or water. Uninoculated controls consisted of 180 µL of M9-glycerol (0.4%) medium, 20 µL control or experimental solution, and 2 µL of 1×PBS.

Bioassay experiments were set up in 96-well microtiter plates (Corning Costar Assay Plate 3904, Bedford, MA), and sealed with air-permeable membranes (Breathe-Easy, Diversified Biotech, Dedham, MA). Cultures were incubated at 37°C for 24 h either in a bench top heated plate shaker (Southwest Science, Roebling, NJ) at 1,200 revolutions per minute (rpm), or a Biotek Synergy 2 plate reader (Agilent, Santa Clara, CA) with linear shaking at 1,140 cpm. OD_600_ values after 24 h were measured in the Biotek Synergy 2 plate reader. Data were plotted and analyzed in Python.

### Cobalamin adsorption assay

Cyanocobalamin stock solutions and dilutions were prepared in sterile ddH_2_O. In triplicate for each concentration and a ddH_2_O control, soil (0.5 g, either unsterilized or autoclaved three times for 30 minutes) was added to a sterile 2.0 mL microcentrifuge tube (VWR, Radnor, PA), along with 1 mL of corresponding corrinoid solution for each sample. Control solutions were also generated in triplicate without soil. Samples were vortexed for 1 minute and incubated at 25°C for 24 h. After incubation, samples were centrifuged for 10 min at 20,187 × rcf, and 90 µL of supernatant removed and placed into a 96-well PCR plate. These samples were incubated at 95°C for 10 minutes, and cooled to room temperature. Samples were quantified using the *E. coli* bioassay as described above.

### Extraction of soil with phosphate buffer

Soil was autoclaved in glass beakers to release corrinoids from cells and the soil matrix (ref. 15). Though corrinoids are stable compounds, because autoclaving has the potential to chemically modify metabolites, we consider our measurements to be lower bounds. For soil autoclaved wet, sterile ddH_2_O (1 mL/g soil) was added prior to autoclaving. Samples were incubated for 30 minutes at room temperature and subjected to steam sterilization on a liquid cycle for 30 minutes. Samples were cooled to room temperature, and steam sterilization was repeated two additional times. Wet soil extract was filtered directly from the supernatant post-sterilization using a 0.2 µm Supor filter (Pall Corporation, Port Washington, NY).

Soil (10 g, fresh or sterilized dry) was added to a 50 mL centrifuge tube (Falcon, Corning, Glendale, AZ). Either sterile ddH_2_O or 0.1 M K_2_HPO_4_, pH 7.0 (25 mL) was added and the samples incubated at 50°C for 30 minutes, and vortexed for 1 minute every 10 minutes. Samples were then centrifuged at 7,197 × rcf for 10 minutes, and supernatants were filtered through a 0.2 µm Supor filter (Pall Corporation, Port Washington, NY). These supernatants were used directly in the bioassay as described above.

### Growth of E. coli bioassay strains in soil

Fresh soil (1 g) was added to a 16×150mm glass test tube. For soil autoclaved wet, sterile ddH_2_O (1 mL) was added. Samples were incubated for 30 minutes at room temperature and subjected to steam sterilization three times as described above.

For experimental samples, M9-glycerol (0.4%) medium (5 mL) containing 1 mM L-Met, 1 nM CN-Cbl, or 1 nM CN_2_-Cbi was added to the sterilized soil as indicated. Samples were then inoculated with 5 µL *E. coli* bioassay strains prepared as above, except that cells were not diluted before inoculation.

Cultures were grown at 37°C at a 30° angle with shaking at 200 rpm for 24 h. For OD_600_ measurements, 200 µL of sample was removed and added to 96-well plates, and absorbance measured as described above. We find, with the use of M9 medium and a 30° tilt of the culture tubes, that soil settles in the test tubes, whereas other media (e.g. LB), faster shaking rates, or steeper angles leads to disturbance of the soil in the medium, preventing facile analysis.

### Determination of the number of extractions required for corrinoid removal

Fresh soil (100 g) and sterile ddH_2_O (100 mL) were added to a 500 mL beaker. Samples were incubated for 30 minutes at room temperature and subjected to steam sterilization on a liquid cycle for 30 minutes. Samples were cooled to room temperature, and steam sterilization was repeated twice. Sterilized soil (3 g) was mixed with M9-glycerol (0.4%) medium (10 mL) in a sterile 15 mL polypropylene centrifuge tube (Corning, Glendale, AZ). Samples were incubated at 50°C for 30 minutes and vortexed every 10 minutes for 1 minute. Samples were then centrifuged at 7,197 × rcf for 10 minutes, and supernatants were filtered through a 0.2 µm Supor filter (Pall Corporation, Port Washington, NY). This process of extraction — addition of fresh medium, incubation with vortexing, centrifugation, and removal — was repeated a total of 12 times. Bioassay strains were prepared as described above, and 2 µL was added to 200 µL of extraction mixture, or to control M9-glycerol (0.4%) medium with additives. Samples were grown in a heated plate shaker (Southwest Science, Roebling, NJ) set to 1,200 rpm at 37°C for 24 h, and the OD_600_ measured. This extraction procedure was scaled up for total corrinoid quantification by performing eight serial extractions on 50 g of soil, resulting in 1 L of soil extract.

### Determination of soil partition coefficient and estimation of soil corrinoid concentration

We use the following set of variables for these calculations:

K_d,corrinoid_ is the partition coefficient for a corrinoid, which is an intrinsic property based upon the soil and the solution used.

[B12_aq,n_] refers to the concentration of corrinoid in the aqueous phase (using ‘B12’ for simplicity) at the nth extraction round, and [B12*_siol,n_*] the concentration of corrinoid in the soil phase in the nth extraction round. Similarly, *vol_soil_* and *vol_aq_* are the volumes of soil or aqueous solution, which are constant between rounds: *vol_aq_* is 0.01 L, and *vol_soil_* is 0.001 L, assuming a soil density of 1 g/mL, similar to soils at this site (ref. 41).

The partition coefficient, *K_d,corrinoid_* is defined as:

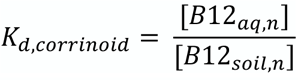

Critical to this analysis is the property that the partition coefficient is the same at each successive round of extraction. So, given an arbitrary extraction round n and n+1:

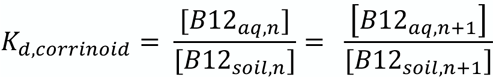

Furthermore, [*B*12_*soil,n*_] is related to the n+1 variables through:

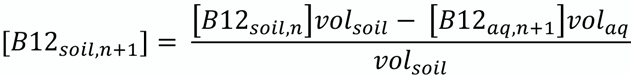

Combining these equations, we obtain:

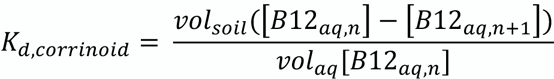

Thus, using extraction rounds 6-9 from Fig. 3A and the bioassay response curve, we can calculate *K_d,corrinoid_* using this formula at three independent stages of the extraction process. Similarly, the total concentration of corrinoid in the soil at an earlier stage is:

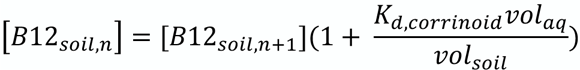

### BtuG expression and purification

We found that further chemical purification of corrinoids from the soil extracts resulted in substantial loss of corrinoid. Therefore, we directly quantified the corrinoids in the soil extract by removing all of the corrinoid present with the high-affinity corrinoid binding protein BtuG (ref. 42), generating a standard curve of the bioassay strain response to added cobalamin in this “corrinoid-free” soil extract, and comparing this response to the extract without corrinoid depletion. BtuG appears to be promiscuous for various corrinoids, binding at fM affinity to both cobalamin and cobinamide (ref. 42). Our results, combined with the fact that corrinoid-dependent growth is not observed in BtuG-treated soil extracts, suggests that this protein is capable of binding to all corrinoids present in soil.

A signal sequence truncation mutant of BtuG2 from *Bacteroides thetaiotaomicron* VPI-5482 with a C-terminal His_10_ tag in a pET21-derived plasmid (ref. 42) was overexpressed in *E. coli* BL21(DE3) Rosetta cells. Briefly, an aliquot of a saturated culture grown in LB supplemented with carbenicillin (100 µg/mL) and chloramphenicol (20 µg/mL) was diluted into LB with the same antibiotics and grown to an OD_600_ of 0.6, after which cultures were induced with 0.5 mM IPTG for 3 h. After centrifugation to isolate the cell pellet, cells were lysed by sonication in lysis buffer (25 mM Tris-HCl (pH 8.2), 500 mM NaCl, 20 mM imidazole, and 5 mM β-mercaptoethanol). Lysate was then clarified by centrifugation at 10,000 × ref for 45 min at 4°C. Clarified lysate was bound to Ni-NTA agarose (Qiagen, Germantown, MD), and the resin was washed three times with 20 mL lysis buffer prior to elution with lysis buffer supplemented with 500 mM imidazole. Purified protein was exchanged into storage buffer (1×PBS, pH 8.0, supplemented with 5% glycerol). The purified protein was concentrated to ∼1 mg/mL by centrifugal filtration (Amicon 30K, Millipore Sigma, Burlington, MA, USA), flash frozen in liquid nitrogen, and stored at −80°C. Protein purity was assessed by SDS-PAGE.

### BtuG-mediated pulldown of corrinoids from soil and quantification of soil corrinoids using corrinoid-depleted soil

To sterile M9-soil extract (10 mL) in a 15-mL centrifuge tube (Olympus, Center Valley, PA) was added 100 µL of either purified BtuG-His_10_ at 17 µM (resulting in a final concentration of 170 nM BtuG-His_10_) or storage buffer (1×PBS, pH 8.0, supplemented with 5% glycerol). Samples were rocked slowly at 25°C for 1 h. Soil extract sample was added to 1 ml Ni-NTA agarose resin (Thermo Scientific, Waltham, MA, US), and samples were incubated with gentle rocking at 25°C for 1 h. Samples were then centrifuged at 2,000 × rcf for 10 minutes to pellet the resin, and supernatants were filtered through 0.22 µm Supor filter (Pall Corporation, Port Washington, NY). Extract solutions treated in this way with BtuG were used as the base medium to generate a standard curve for the bioassay strain’s response to soil extract devoid of corrinoids, whereas the soil extract that was not treated with BtuG was used to measure the corrinoid content of the soil extract.

### Guided biosynthesis and purification of non-commercially available corrinoids

Non-commercially available corrinoids were produced in bacterial cultures, extracted, and purified as described (ref. 40, 43). Specifically, [Ade]Cba and [2-MeAde]Cba were purified from cultures of *Propionibacterium acidipropionici* strain DSM 20273, [Cre]Cba from *Sporomusa ovata* DSM 2662, and [5-OHBza]Cba from *Propionibacterium freudenreichii* CICC 10019 as previously described (ref. 44), using fermentation medium supplemented with 5-hydroxybenzimidazole at 10 mg/L. We acknowledge the possibility that low abundance non-UV-active compounds copurifying with these corrinoids may play additional roles in altering the soil communities.

### Establishing soil-derived enrichment cultures with varying corrinoids

Inocula for soil-derived enrichment cultures were generated from soil (2.5 g) resuspended in sterile 1×PBS (25 mL) supplemented with 2.24 mM sodium pyrophosphate (Alfa Aesar, Heysham, England) and stirred for 30 minutes. The soil slurry was allowed to settle for 15 minutes before use in generating enrichment cultures. 1 mL cultures containing M9 or R2 medium (prepared as in Supp. Table 2) supplemented with 10 nM corrinoid, or no corrinoid, and inoculated with 20 µL of soil suspension were grown in 96-well deep-well plates (Greiner, Monroe, NC) with six biological replicates per condition plus one uninoculated sample. These enrichment cultures were incubated in the dark at 25°C, without shaking. In all inoculated samples, a clear biofilm was observed at the bottom of wells after one week, whereas no biofilm was observed in uninoculated controls. Samples were passaged into fresh medium at a 1:11 dilution weekly, by transferring 100 µL from each culture into 1 mL fresh medium, without disturbing the biofilm forming at the bottom of each well. The remaining sample was harvested by centrifugation at 4,000 × ref for 3 minutes and stored at −80°C for DNA extraction.

### Extraction of DNA from enrichment cultures

Total DNA from enrichment cultures was extracted using a method described in (ref. 45). Briefly, samples were resuspended in 100 µL of a 1% IGEPAL CA-630 (Sigma, Burlington, MA) solution, obtained by dissolving 1 mL of IGEPAL into 99 mL sterile water, transferred to PCR plates (GeneMate, Milford, Surrey), and sealed with foil film. Samples were frozen at −20°C and subsequently thawed and brought to room temperature a total of five times. To each sample, Proteinase K was added to a final concentration of 100 µg/mL (Thermo Scientific, Waltham, MA). Samples were incubated at 60°C for 1 h in a thermocycler followed by 95°C for 15 minutes to inactivate Proteinase K.

### Direct corrinoid additions to soil microcosms and DNA extractions

We established 155 1-g soil microcosms containing 22 µL of water or a 10 nM solution of one of six corrinoids (Fig. 1A) in water, resulting in a final corrinoid concentration of 220 pmol/g soil, with five replicates of each condition and time points except time 0, which had five replicates in total. A set of microcosms with no added water or corrinoid was also included for time 0. Soil microcosms were destructively harvested such that after 0, 3, 10, 30, and 50 days microcosms were transferred to −80°C for storage. For all DNA extractions from soil we used the DNeasy PowerSoil Pro Kit (250) (Qiagen, Germantown, MD).

### Community composition assessed by 16S rRNA gene amplicon sequencing

The 16S rRNA gene V4/V5 region was amplified by PCR and analyzed as previously described in (ref. 46). Data for enrichment and microcosm relative abundances are found in Supp. Tables 3 and 4. We observe some cross-contamination in our enrichment extractions, as seen by uninoculated controls containing reads in comparable amounts to the inoculated samples, as has been observed in other 16S rRNA gene surveys (ref. 47, 48). Given that we observed no growth in these uninoculated samples, we expect that these reads are representative of minor cross-contamination between wells during the extraction process. However, as a secondary check we performed ordination analysis between the inoculated and uninoculated controls at each week using both Jaccard and Bray-Curtis metrics, and observed significant differences between the cluster of inoculated samples and cluster of uninoculated controls (*P* ≤ 0.05 in all cases according to PERMANOVA). Furthermore, we see minimal reads for the genera that possess significant differences between corrinoid conditions (*Rhizobium*, *Shinella*, and *Microbacterium*), and we observe 1,195 distinct zOTUs: 1,022 solely observed within enrichment samples, 45 solely within uninoculated samples, and 128 in both. As such, the actual corrinoid-dependent changes in these enrichments are likely to be larger than what we observed.

### Ordinations and data visualization

All analyses were performed using Python 3.8.2 (docs.python.org/3.8/) and pandas version 1.5.1 (ref. 49, 50). Graphs were generated using seaborn version 0.11.0 (ref. 51). We constructed ordination plots to analyze compositional differences across time and corrinoid treatment. We constructed a matrix of zOTUs and their frequency per sample (corrinoid added, time point, replicate) by iterating through each taxonomic level from genus to phylum and aggregating the frequency of all zOTUs that fell within subset phylogenetic classifications. From these matrices of frequency per taxon per sample, we used the python scikit package to calculate Bray-Curtis dissimilarity and to perform principal coordinates analysis (PcoA) to visualize the dissimilarity between samples. To test corrinoid treatment significance we ran ANOSIM and PERMANOVA tests using the scikit-bio Python package on Bray-Curtis distance matrices from our 16S rRNA gene amplicon sequencing data.

### Identifying soil taxa sensitive to corrinoid treatment

To identify taxa that showed significantly different frequencies across corrinoid conditions, we iterated through phylogenetic levels of classification and ran Kruskal-Wallis tests using the scipy statistics package (ref. 52) across replicates per corrinoid condition per time point. For analysis of enrichments, we identified the fraction of taxa sensitive to corrinoids at each taxonomic level by dividing the set of all taxa at each taxonomic level that were corrinoid-responsive at any time point by the set of all taxa observed at that level. We visualized these data as bar plots per taxonomic level using seaborn which aggregated all timepoints and media conditions to display the total fraction of taxa sensitive to corrinoids in all enrichments. For analysis of soil microcosms, we summed the frequency of Kruskal-Wallis statistically significant (*P* value ≤ 0.05) taxa at each phylogenetic level (phylum through zOTU) to quantify the proportion of taxons detected from soil microcosms that were responsive to different corrinoid types (on average across corrinoid conditions). We visualized these data as bar plots through time using seaborn. We plotted the four phyla – *Actinobacteria*, *Firmicutes*, *Bacteroidetes*, *Verrucomicrobia* – that showed statistically significant distinct responses to varying corrinoid types also as bar plots using seaborn. We generated the same data visualization for the order that showed the highest frequency and statistically significant response to corrinoids, *Actinomycetales*, and the highest frequency statistically significant zOTU identified as *Arthrobacter psychrochitiniphilus*. We determined pairwise statistically significant comparisons between corrinoid treatments using the Mann-Whitney test.

## Results

### Genome-resolved metagenomics reveal widespread corrinoid dependence in soil

We investigated whether corrinoid-producing and corrinoid-dependent microbes (Fig. 1B) coexist in soil from the Northern California grassland field site at Hopland Research and Extension Center by assessing the capacity of 503 MAGs (ref. 36, 37, 38) to produce and use corrinoids. These MAGs encompass 10 bacterial and one archaeal phyla, and are predominantly *Proteobacteria* (31%) and *Actinobacteria* (23%) which represent the abundant phyla at this field site (ref. 53, 54).

First, we sought to understand how soil microbes use corrinoids. In both archaeal and bacterial MAGs, we found that the most prevalent corrinoid-dependent enzymes are methionine synthase (MetH), responsible for the last step in methionine production (ref. 55), and methylmalonyl-CoA mutase (MCM), which is involved in propionate and branched-chain amino acid metabolism (Fig. 2A) (ref. 33). The majority of archaeal MAGs additionally encode corrinoid-dependent methyltransferases, used in methanogenesis (ref. 56) (Fig. 2A). The four most prevalent corrinoid-dependent functions in bacterial MAGs are those with known corrinoid-independent alternatives: the corrinoid-independent methionine synthase (ref. 57), the methylcitrate pathway as an alternative to MCM (ref. 58), the corrinoid-independent epoxyqueuosine reductase that modifies tRNAs (ref. 59), and corrinoid-independent ribonucleotide reductases (RNRs) involved in deoxyribonucleotide synthesis (ref. 60) (Fig. 2B). Corrinoid-independent MCM and RNR are slightly more prevalent in these MAGs than their corrinoid-dependent counterparts: 22% of MAGs contained only the corrinoid-independent methylcitrate pathway (MCM alternative) and 55% of MAGs with a detected RNR maintain only the corrinoid-independent alternative. In contrast, 61% of the MAGs encode the corrinoid-dependent epoxyqueuosine reductase, although only one encodes the corrinoid-independent version, suggesting corrinoids are required for this function in soil. A substantial proportion of MAGs contained functionally redundant corrinoid-dependent and -independent enzymes and pathways. Specifically, 43% of the MAGs that encode the corrinoid-dependent methionine synthase also contained the corrinoid-independent version, and 9% of MAGs contained both a corrinoid-dependent and -independent RNR (Fig. 2B). Thus, a subset of soil microbes may switch between corrinoid-dependent and -independent processes, presumably based on environmental corrinoid availability (ref. 61, 62).

**Figure 2.**
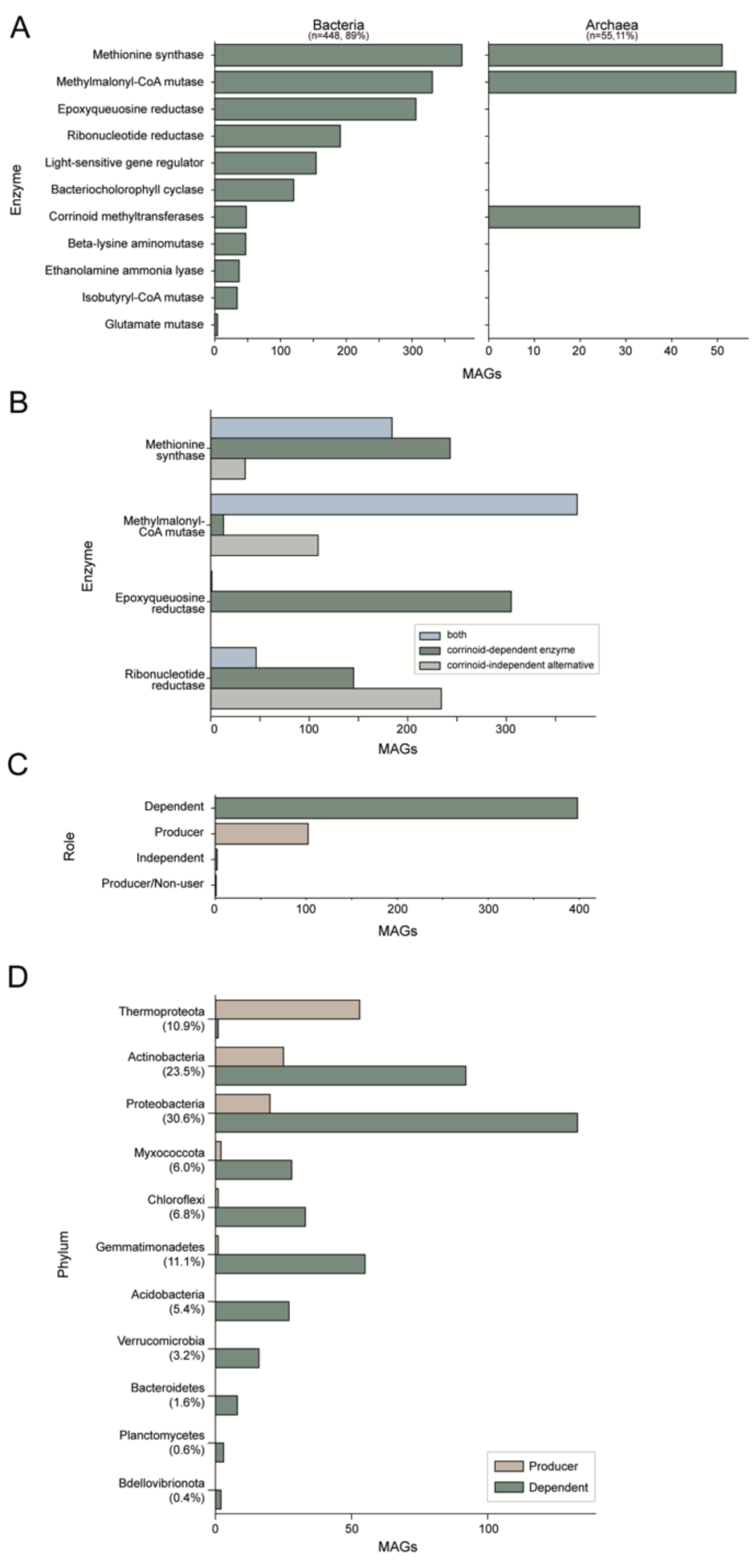
Analysis of corrinoid production and dependence in soil MAGs. A. Corrinoid-dependent processes encoded in soil MAGs split by bacteria (left) and archaea (right). B. The number of bacterial and archaeal MAGs that encode corrinoid-dependent enzymes, corrinoid-independent alternatives, or both are shown for the four most abundant corrinoid-dependent processes in soil. C. The number of MAGs categorized by corrinoid ecological role (dependent, producer, independent, and producer/non-user) is shown for the complete dataset. D. The number of corrinoid producer and dependent MAGs classified by their phyla. The proportional representation of each phylum of the 503 MAGs assessed is shown as a percentage under each phylum name on the y axis label.

We assessed the extent to which corrinoid-based interactions may occur by predicting which MAGs represent predicted corrinoid “producers” – capable of producing corrinoids for their metabolism – or “dependents” – reliant on external corrinoids for corrinoid-dependent processes. In addition to these ecological roles, we expected some organisms to be “independents”, lacking pathways for producing or using corrinoids (Fig. 1B). Each MAG was assigned a corrinoid metabolism category based on the presence or absence of corrinoid-dependent enzymes and whether they contained at least one of six marker genes that are 99% correlated with *de novo* corrinoid biosynthesis in reference bacterial genomes (ref. 16). The majority of MAGs (79%) were predicted dependents, whereas 20% were classified as producers and less than 1% (two MAGs) as independents (Fig. 2C). However, we found that one archaeal MAG of the genus *Nitrososphaera* could be classified as a “producer/non-user,” predicted to be capable of synthesizing corrinoids but lacking corrinoid-dependent enzymes (ref. 16). Though the producer/non-user category has been previously identified in MAGs (ref. 22), it has not been previously identified in complete genomes (ref. 16), and we could not verify this predicted classification. Additionally, from a concurrent laboratory experiment that used stable isotope probing via addition of ^18^O-labeled water (H_218_O) to identify actively replicating microbes (ref. 37), 72 of these annotated MAGs – nine predicted producers and 63 dependents – were detected as active within one week after H_218_O addition. Thus, corrinoid-producing and - dependent microbes are not only present, but are active members of the soil microbiome.

We found that the predicted capacity to produce and use external corrinoids is not evenly distributed across phyla in this set of MAGs. Most predicted corrinoid producers were *Thermoproteota*, and in this lineage, all except one MAG were predicted to be able to synthesize a corrinoid (Fig. 2D). *Actinobacteria* and *Proteobacteria* had the highest number of dependents overall and together contained nearly half (44%) of all producers. The remaining phyla had few or no producers (Fig. 2D). These results suggest that corrinoid production is phylogenetically unevenly distributed in this soil, with *Thermoproteota*, *Actinobacteria*, and *Proteobacteria* serving as the major suppliers of corrinoids.

### Soil corrinoids are abundant and adhere to the soil matrix

The finding that the vast majority of microbes in Hopland soil encode corrinoid-dependent processes, yet cannot produce corrinoids, suggests that corrinoids are shared nutrients in this community. We therefore sought to quantify the corrinoids in this soil to determine the extent to which corrinoids are limiting nutrients for microbes in this environment. Because of the physical and chemical complexity of soil, chemical analysis methods such as LC-MS require multiple purification steps that reduce yield and obscure quantification (ref. 63). Thus, we measured the total corrinoid in soil samples using an *E. coli* bioassay that can accurately measure corrinoid levels in crude extracts (ref. 40). This bioassay is broadly responsive to distinct corrinoids, although with different sensitivities; thus we report our results in terms of cobalamin equivalents. To prepare samples for analysis, we first used a previously applied method for organic extraction of corrinoids from a similar soil (ref. 21). Despite the wealth of genomic data pinpointing corrinoids as important in this soil, we detected little to no corrinoid in these extracts or in extractions using ten other organic solvent mixtures (ref. 64) (Supp. Fig. 1). Further, even when soil was spiked with solutions containing up to 1.5 nM cobalamin, no corrinoid was detected in extracts, despite the fact that the bioassay response saturates at 1 nM cobalamin (Supp. Fig. 2). These results suggest that the partition coefficient for corrinoids (a measure of relative solubility) under these solvents is less than 5×10^-5^ (see Methods). Our results suggested that soil adsorbs corrinoid, and the adsorbed corrinoid cannot be recovered effectively by organic extraction.

Unlike in extractions with organic solvents, we detected corrinoids after soil was autoclaved and subsequently extracted with a phosphate-buffered solution. Both steam sterilization and phosphate contributed to the extraction efficiency (Supp. Fig. 3), possibly by facilitating corrinoid release from cells and macromolecules by heat and phosphate-mediated nonspecific anion exchange interactions, respectively (ref. 65). The bioassay response was diminished to a baseline level after eight successive extractions with M9 medium (a medium containing 60 mM phosphate) from the same soil (Fig. 3A, Supp. Fig. 4). Using the change in bioassay response from successive extractions, we calculated a partition coefficient for corrinoid of 4×10^-2^ ± 8×10^-3^ under these conditions. The extracted product appeared to represent most of the corrinoid in the sample, as the post-extraction soil showed a low corrinoid signal (Fig. 3B). Using this partition coefficient combined with the corrinoid quantified those extraction steps with bioassay responses in the linear quantitation range, we calculate that the soil initially contained 34 ± 4 pmol cobalamin equivalents per gram. In agreement with this extrapolated measurement, quantification of the pooled extracts showed that the soil contained 41 pmol cobalamin equivalents per gram (Supp. Fig. 5), though current methods do not allow us to distinguish intracellular from extracellular corrinoids. Combined with our previous LC-MS results surveying the corrinoid diversity of this soil (ref. 66), we expect that the majority of this corrinoid reservoir is cobalamin. Based on an estimate of 10^9^ microbial cells per gram of soil, this soil contains approximately 10^4^ corrinoid molecules per microbial cell, 1-2 orders of magnitude above corrinoid concentrations previously determined to support maximal growth of bacteria in pure culture, and far above that required by corrinoid-dependent isolates obtained from this soil (ref. 14, 18, 67, 68). However, given that cobalamin strongly adsorbs to soil and that the ratio of intracellular to extracellular corrinoid is unknown, the extent to which these corrinoids are accessible to microbes that require them remains unclear.

**Figure 3.**
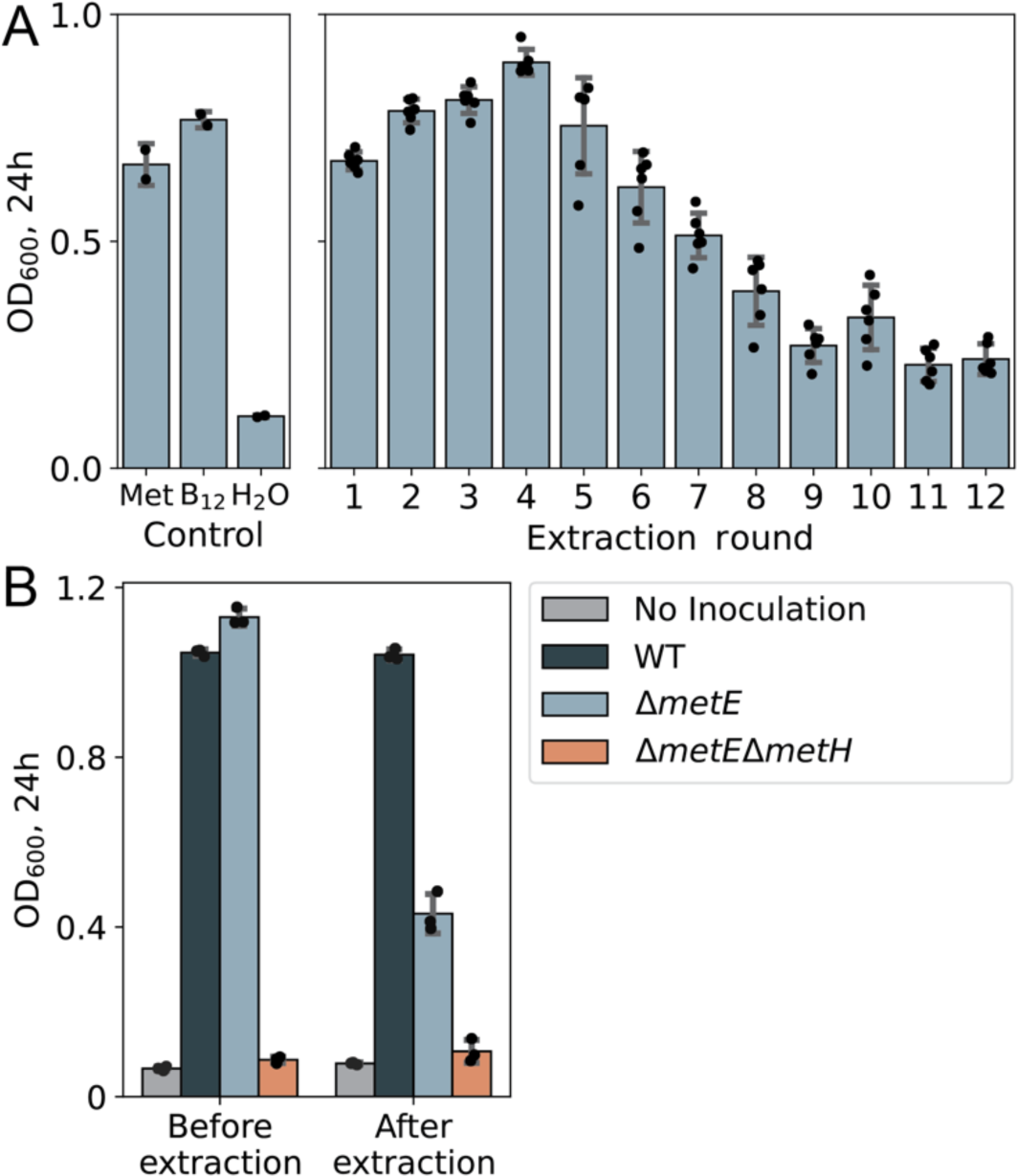
Quantification of total corrinoid in M9 soil extracts by microbiological assay. A. Growth (measured as OD600) of corrinoid-responsive *E. coli ΔmetE* strain in M9-glycerol medium with _L_-methionine (Met, 6.7 mM), cyanocobalamin (B12, 1 nM), or no additional supplement (H2O), or in serial extracts from soil obtained using M9-glycerol. B. OD600 of the corrinoid-responsive *E. coli ΔmetE* and control strains in M9-glycerol incubated with steam-sterilized untreated soil (“Before extraction”) or after the 12^th^ round of corrinoid extraction (“After extraction”).

### Corrinoid structure influences community assembly in soil-derived enrichment cultures

Having established that corrinoid metabolism is prevalent in soil microbes and that corrinoids are abundant in this soil, we examined whether corrinoids that vary in lower ligand structure – known to impact the growth of individual microbes (ref. 14, 33) – can elicit changes at the community level. To that end, we established soil-derived enrichment cultures containing a panel of in-house extracted and purified corrinoids. We seeded two basal media (M9 and R2) with a soil suspension and added one of six corrinoids (Fig. 1A) or no corrinoid. We diluted the cultures 11-fold into fresh medium every 7 days and analyzed community profiles by 16S rRNA gene amplicon sequencing at stages previously shown to be ‘early’ (1 and 2 weeks) and ‘late’ (12 and 14 weeks) for community assembly and stabilization (ref. 69). As observed previously with similar enrichments, the 16S rRNA gene composition showed greater consistency at higher taxonomic levels: by week 14, six of the 12 orders observed at week 1 were present in 95% of enrichment cultures. In contrast, of the 378 total zOTUs, only one was present in a majority of the cultures at week 14 (ref. 69).

The presence of corrinoids in soil-derived enrichments impacted microbiome composition at all taxonomic levels (Kruskal-Wallis, *P* ≤ 0.05). Quantifying the total number of taxa responsive to corrinoid addition across both media and time, we found over 20% of all orders detected from the 84 soil-derived enrichment cultures responded to corrinoid addition (Fig. 4A). In the defined minimal M9 medium, corrinoid amendment led to divergent taxonomic compositions that were most pronounced at week 2, as measured by both Bray-Curtis and Jaccard dissimilarity indices (Fig. 4B, Supp. Figs. 6-8). As observed in a previous study of soil enrichment cultures, replicate enrichments separated into two distinct community types, each dominated by members of a different order (ref. 69). In contrast, enrichment cultures seeded in a modified version of the complex R2 medium did not cluster by corrinoid condition (*P* > 0.05) (Fig. 4C, Supp. Figs. 6-8). Comparing across media conditions revealed that the effect size of corrinoid treatment is comparable to that of the different basal media (By ANOSIM, *r* = 0.0652, *P* ≤ 0.05 when clustered by medium, compared to *r* = 0.1330, *P* ≤ 0.05 when clustered by corrinoid) (Supp. Fig. 9), demonstrating that a single change to corrinoid structure was as impactful to microbiome composition as the nutritional differences between defined and complex media.

**Figure 4.**
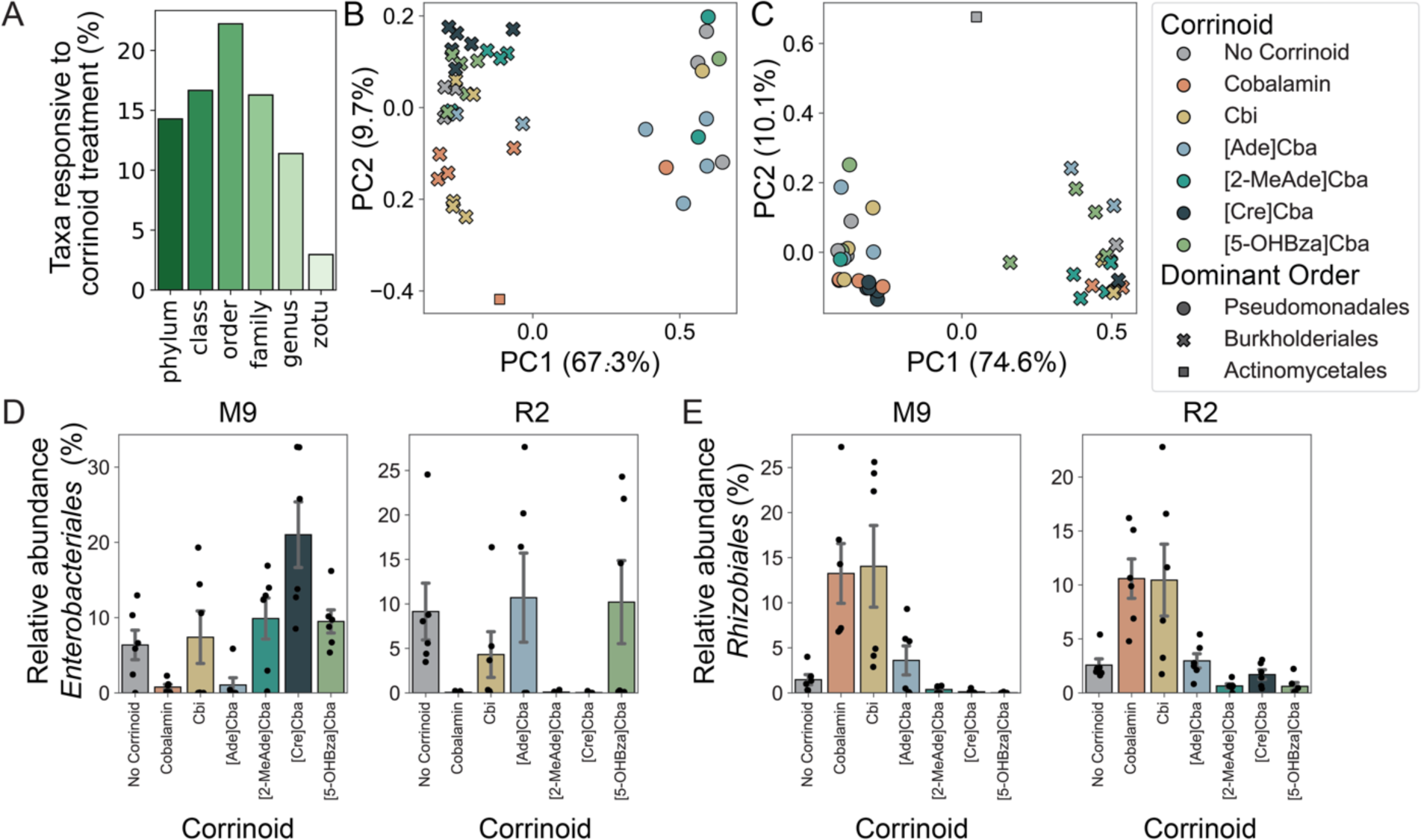
Responses of soil-derived enrichment cultures to corrinoids. A. For each taxonomic level, the fraction of all taxa observed in enrichment cultures that are corrinoid responsive (Kruskal-Wallis, *P* ≤ 0.05) at any timepoint. B, C. Principal coordinate analysis (PCoA) of Bray-Curtis dissimilarity for order-level taxonomy of enrichments passaged in M9 (B) and R2 (C) media, grouped by corrinoid condition at 2 weeks. M9 enrichments (B) significantly cluster by corrinoid (ANOSIM: *r* = 0.13, *P* ≤ 0.05; PERMANOVA: *r* = 2.13, *P* ≤ 0.05). R2 enrichment cultures (C) do not cluster by corrinoid (*P* > 0.05). D,E. Relative abundance of *Enterobacteriales* (D) and *Rhizobiales* (E) in M9- or R2-derived enrichment cultures at week 2. Error bars represent standard error of the mean.

Further analysis showed that, for both media formulations, specific taxa were corrinoid-responsive, and that the profile of these responses were dependent upon both the corrinoid structure and medium used (Fig. 4D,E). The impact of corrinoid amendment on the *Enterobacteriales* order (represented by only one zOTU) appeared to be influenced by the medium used (Fig. 4D). In contrast, the *Rhizobiales* order (represented by three zOTUs) showed similar sensitivities across corrinoid conditions irrespective of differences in the growth medium or community context, suggesting its response was directly influenced by the structure of the added corrinoid (Fig. 4E). Furthermore, these impacts were corrinoid-specific: in most corrinoid-responsive taxa, corrinoid addition led to increased abundance, although we observed that *Enterobacteriales* relative abundance was suppressed by multiple corrinoids (Fig. 4D). These results demonstrate that corrinoid structure can alter the abundances of specific taxa, which could have cascading community-wide effects, and further suggests that some microbes may be directly influenced by added corrinoids whereas others may be indirectly affected.

### Corrinoids undetected in soil transiently influence the composition of soil communities

We investigated whether *in situ* corrinoid amendment could influence the composition of established soil communities. We amended soil from the same research site as that used for corrinoid quantifications and establishment of soil-derived enrichments with the same seven corrinoid conditions used in the enrichments, destructively harvested samples after 0, 3, 10, 30, and 50 days, and performed 16S rRNA gene amplicon sequencing to determine community composition. These microcosms were dominated by *Actinobacteria* and *Proteobacteria* (Supp. Fig. 10), aligning with our MAG and previous data from this site as the most abundant bacterial phyla present (ref. 36, 38, 53).

Consistent with our hypothesis that corrinoids influence community structure, we found that corrinoid addition impacted the taxonomic composition of soil microcosms. Corrinoid treatment significantly altered a percentage of the total community ranging from approximately 70% at the class level to 19% at the zOTU level just three days after treatment (Fig. 5A). The percentage of taxa significantly sensitive to corrinoids (Kruskal-Wallis test, *P* ≤ 0.05) decreased through time on all phylogenetic levels (Fig. 5A). Further, we observed distinct effects of different corrinoids on community composition. Principal coordinate analysis of Bray-Curtis dissimilarity matrices of relative abundances by order showed significant differences (*P* ≤ 0.05, ANOSIM or PERMANOVA) across corrinoid conditions three days after treatment, with soil amended with cobalamin clustering most closely with unamended soil. Addition of [2-MeAde]Cba, [Cre]Cba, [Ade]Cba, and [5-OHBza]Cba, and to a lesser extent, Cbi, showed profiles distinct from unamended soil and the cobalamin treatment (Fig. 5B). The treatment effect observed at day 3 was largely lost by day 50 (Fig. 5B, Supp. Fig. 11), suggesting that one-time corrinoid addition is a significant disturbance from which the community can rebound. Together, these results demonstrate that addition of corrinoids other than cobalamin – the most prevalent corrinoid detected in this soil – significantly, but transiently, impacts soil community composition.

**Figure 5.**
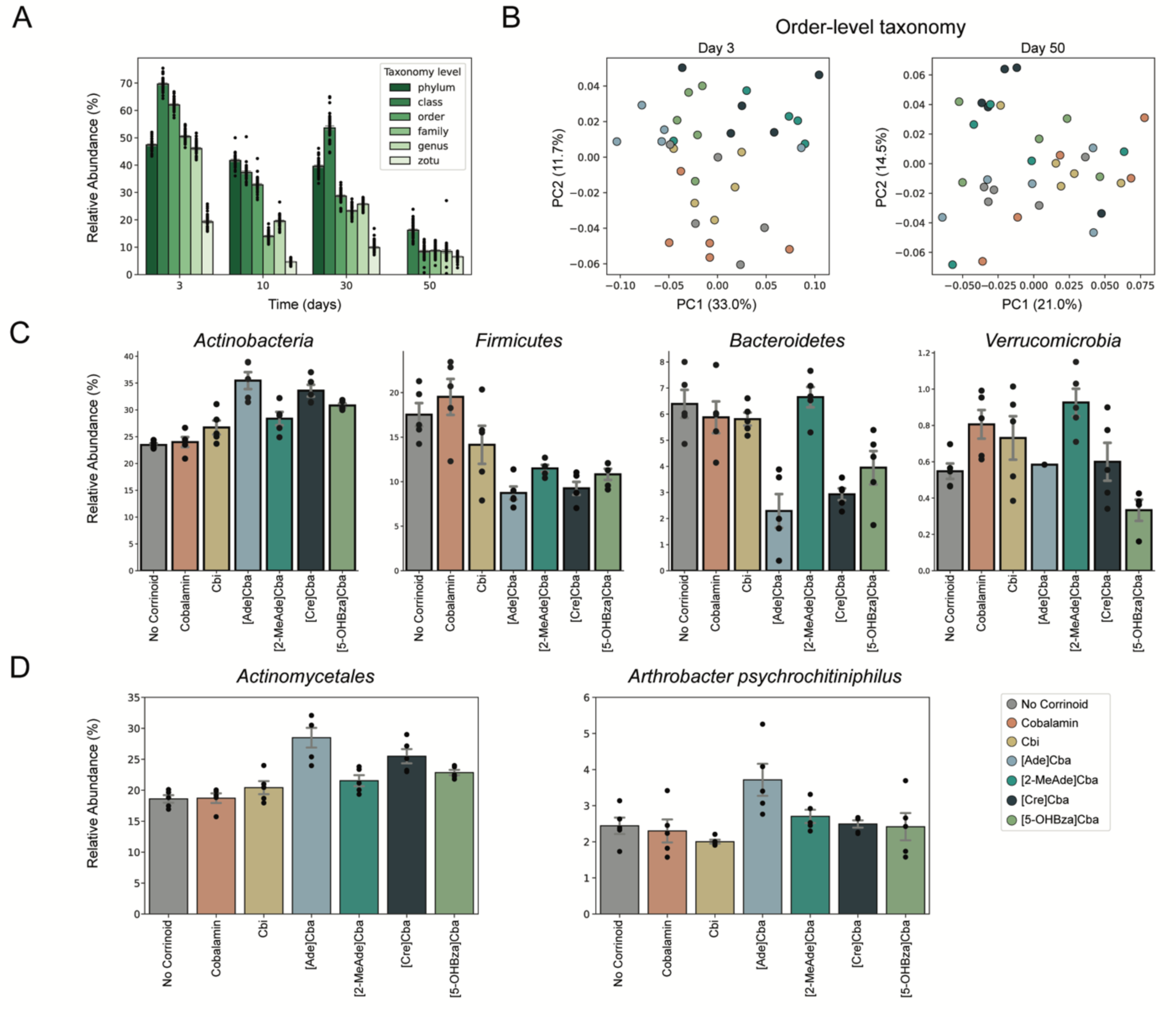
Treatment effect of corrinoids on soil microcosms. A. The proportion of the total community at each taxonomic level (shades of green) that were statistically significantly (Kruskal-Wallis, *P* ≤ 0.05) sensitive to corrinoid treatment through time. B. Principal coordinate analysis of Bray-Curtis dissimilarity for order-level taxonomy on day 3 (left) and day 50 (right), grouped by corrinoid condition. Each plot shows the first two principal coordinates with the percent variance explained labeled on each axis. Microcosms significantly cluster by corrinoid at day 3 (ANOSIM: *r* = 0.21, *P* ≤ 0.05; PERMANOVA: *r* = 1.95, *P* ≤ 0.05), but not at day 50 (ANOSIM, PERMANOVA *P* ≥ 0.05). C. Relative abundances in each treatment for phyla with statistically significant responses to corrinoid amendment at day 3. D. Relative abundances of the most prevalent *Actinobacteria* order, *Actinomycetales*, and the most abundant zOTU, *Arthrobacter psychrochitiniphilus*, which responded to corrinoid. Error bars represent standard error of the mean across replicates. Underlying points show five replicates per treatment per time point.

To determine which taxa underlay the compositional differences we observed among corrinoid treatments, we identified taxa with statistically significant differences (Kruskal-Wallis test, *P* ≤ 0.05) in relative abundance across corrinoid amendment conditions on each phylogenetic level at day 3. At the phylum level, we observed significant differences in relative abundance for *Actinobacteria*, *Firmicutes*, *Bacteroidetes*, and *Verrucomicrobia* (Fig. 5C). Each of these phyla showed a unique pattern of relative abundances across corrinoid conditions. For *Actinobacteria* and *Firmicutes*, abundances were similar in the no corrinoid and cobalamin conditions, and were significantly distinct (Mann-Whitney, *P* ≤ 0.05) from [2-MeAde]Cba, [Cre]Cba, and [5-OHBza]Cba, with *Actinobacteria* enhanced and *Firmicutes* relatively less abundant in the latter corrinoids. Like *Firmicutes*, *Bacteroidetes* were significantly elevated in relative abundance in the no corrinoid and cobalamin conditions, but were also enhanced in Cbi and [2-MeAde]Cba. In contrast, *Verrucomicrobia* exhibited a significant difference in relative abundance with cobalamin addition compared to the no corrinoid treatment. The most prevalent phylum in these samples, *Proteobacteria*, represented 40.7% of the community at day 3, but showed significant treatment effects only at day 10 (Supp. Fig. 11).

Within the phylum *Actinobacteria*, the order *Actinomycetales* comprised the majority of sequences and showed trends similar to the *Actinobacteria* phylum as a whole. Specifically, addition of cobalamin did not significantly influence *Actinomycetales* abundances at 3 days, but addition of [Ade]Cba, and to a lesser extent, [Cre]Cba, led to a significantly higher abundance compared to all other corrinoids (Mann-Whitney, *P* ≤ 0.05). A single *Actinomycetales* zOTU, identified as *Arthrobacter psychrochitiniphilus*, was significantly impacted by [Ade]Cba addition. This zOTU represented approximately 3.7 ± 0.4% of the total community in this treatment (Fig. 5D), compared to 2.4 ± 0.1% in all other treatments (Kruskal-Wallis test, *P* = 0.024). This example illustrates how a single zOTU can explain a portion of the differences observed at higher taxonomic levels. Together, these results demonstrate that soil community composition can be substantially and distinctly altered by addition of different corrinoids, with community changes driven by distinct responses of specific taxonomic groups.

## Discussion

A growing body of work has demonstrated that the identity and availability of nutrients influences function in soil microbial communities (ref. 4, 70, 71, 72), yet the impact of shared micronutrients on soil microbiome structure remains unknown. Corrinoids are an ideal model to understand how microbial communities respond to micronutrients: the ecological role of bacteria and archaea with respect to corrinoids can be readily predicted by specific genomic signatures for corrinoid biosynthesis and use, and the preferences of individual microbes for specific corrinoids suggest that distinct corrinoids could target subsets of a complex community. In this study, we applied the corrinoid model to soil, considered one of the most complex microbial ecosystems. We used metagenomic sequencing to classify the ecological roles that members of the microbiome perform with respect to corrinoid metabolism, quantified the corrinoid composition and content of a grassland soil, and measured the compositional shifts in the microbiome in response to structurally distinct corrinoids in both soil and soil-derived enrichment cultures.

We previously found (ref. 66), in agreement with other studies of corrinoids in soils (ref. 21), that the most prevalent corrinoid in soil is cobalamin. Two prior studies that quantified corrinoids from several soils used different extraction techniques (ref. 20, 21). The sampled soils in both studies exhibited up to 50-fold differences in the amount of total corrinoid found across different soils. However, the two extraction methods differed in the average corrinoid content found in soils by an order of magnitude, suggesting different extraction efficiencies for the two methods. In the present study, we developed an extraction method that differs substantially from common metabolite extraction procedures to iteratively remove corrinoids from soil to near-depletion. We reveal that our soil environment contains a deep reservoir of corrinoids, of which the majority is cobalamin (ref. 66), that exceeds the minimum corrinoid requirements determined for individual bacteria in culture (25-500 molecules/cell (ref. 67, 68), compared to approximately 10^4^ molecules/cell detected in soil). The high concentration of corrinoids in soil could support the maintenance of widespread corrinoid-dependent metabolism observed in MAGs. Conversely, the prevalence of corrinoid-independent alternative enzymes may indicate inconsistent corrinoid availability, especially compared to marine systems where genomes show streamlining and loss of corrinoid-independent alternative enzymes (ref. 73). Further, the requirements for steam sterilization and serial extractions to remove corrinoids from the soil matrix suggest that, although likely present in excess, much of the corrinoid in soil could be inaccessible to the microbes that require them, due to sequestration by either the soil matrix or by other microbes. Natural fluctuations in precipitation and temperature may influence the accessibility of soil corrinoids, suggesting that the soil corrinoid reservoir, as in marine systems, may be seasonally dynamic (ref. 17). Further, anthropogenic inputs such as phosphate-containing fertilizer, analogous to the high phosphate content in our extraction solution, could influence corrinoid availability (ref. 74). Our emerging understanding of the chemistry of corrinoids in soil aligns with other nutritional patterns of environment-influenced adsorption and release from the soil matrix (ref. 71, 75). The influence of environmental factors on the availability of corrinoids and other nutrients likely has downstream effects on soil community structure and function.

We propose two, non-mutually exclusive hypotheses for the mechanism of corrinoid cycling in soil (Fig. 6). First, corrinoids may be released by producers and directly taken up by dependent organisms, as observed for microbes co-cultured in liquid media (ref. 76, 77). Alternatively, the soil matrix may serve as a primary medium of this cycling process, where environmental factors such as soil water content mediate the propensity of corrinoids released by producers to become adsorbed to the soil matrix versus becoming accessible to dependent organisms. In this scenario, the ability of solution or can use matrix-associated corrinoids. Given our microcosm results, we expect that microbes have developed strategies to liberate these matrix-associated corrinoids, analogous to solubilization of phosphate from soil (ref. 78). In addition, because the mechanism of corrinoid release from producer cells remains unknown, it is ntify from genome sequences ir original source would lead to.

**Figure 6.**
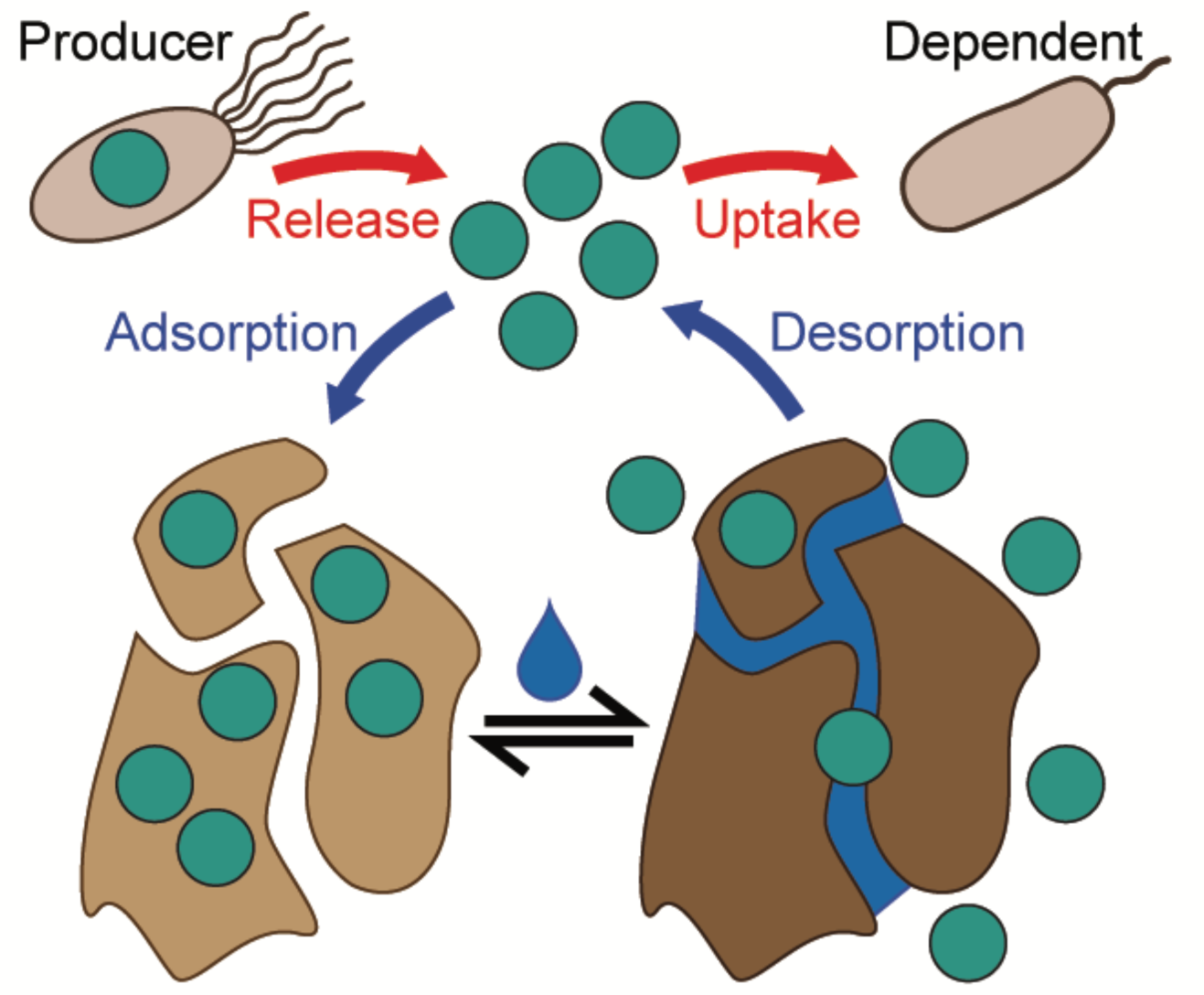
Model for corrinoid interactions in soil. Corrinoid producers synthesize corrinoids, which are released through unknown mechanisms. A portion of this soluble corrinoid pool adsorbs to soil and may become mobilized, potentially via soil wetting or in response to other nutritional, environmental, or anthropogenic inputs. Corrinoids present in the soluble portion of this reservoir may be taken up by corrinoid-dependent microbes following release by producers or desorption from the soil matrix.

The abundance of cobalamin in soil may explain the discrepancy in the effect of cobalamin addition between soil-derived enrichment cultures and soil microcosms (Figs. 3, 4). We found that addition of cobalamin substantially impacted community composition in the enrichments but showed a minimal impact on soil microcosms, whereas addition of other corrinoids had profound and significant impacts in both enrichments and soil. These results could mean that much of the cobalamin detected in soil is bioavailable, and that cobalamin is not a limiting micronutrient in this environment. In nature, the introduction of corrinoids other than cobalamin to soil may mimic the interplay between soil fauna and microbes; the gut and fecal contents of several invertebrates and macrofauna have been found to contain diverse corrinoids that microbes may transiently encounter (ref. 66). Indeed, we have observed that corrinoid-dependent taxa isolated from this soil are able to use many corrinoids undetected in this soil, with different affinities for each (ref. 18).

We observed that corrinoids with similar chemical structures did not promote similar community-wide compositional changes. The two purinyl ([Ade]Cba and [2-MeAde]Cba) compared to the two benzimidazoloyl corrinoids (cobalamin and [5-OHBza]Cba) resulted in distinct community compositions both in soil microcosms and enrichments. Corrinoids within the same structural class can exhibit different effects both on individual enzymes (ref. 33) and whole organisms (ref. 14), suggesting agreement between studies at the molecular and community scales. Our metagenomics analysis suggests that the key corrinoid-dependent enzymatic processes that would be impacted by corrinoid addition are methionine synthesis (MetH), carbon cycling through propionate (MCM), and queuosine biosynthesis (QueG). We expect that the direct effects of these corrinoid-dependent processes on the growth of individual microbes yield additional indirect effects, where microbes respond to the corrinoid-mediated growth enhancement or inhibition of other microbes. Alterations in carbon flux caused by MCM activity or stress responses mediated by QueG are potential mechanisms by which these indirect effects are mediated. Specifically, we attribute the differential corrinoid response of the *Enterobacter* zOTU in the different enrichment media to indirect effects (Fig. 4D): for instance, R2 contains methionine from tryptone, which could circumvent an organism’s requirement for corrinoid-dependent methionine synthase activity. In contrast, the similar corrinoid responses of *Rhizobiales* observed in different media may be due to a direct response to corrinoids (Fig. 4E). In soil – a complex substrate inhabited by a diverse community – a combination of these direct and indirect effects would lead to significant and detectable changes in the community upon nutrient addition.

We find that, although different corrinoids cause profound restructuring of both microcosm and enrichment communities, these corrinoid-dependent differences in community composition are transient. The soil community appeared initially sensitive to corrinoid addition and ultimately rebounded, analogous to the resilience of gut microbial communities to short-term antibiotic use (ref. 80). The outcome of the added corrinoids remains unclear; they may be adsorbed by the soil and become inaccessible, transformed into other corrinoids by a process known as remodeling, or degraded by light or other yet-unknown processes (ref. 29, 32, 81, 82, 83, 84).

From a broader perspective, our development of the corrinoid model in soil provides an important case study to investigate the response of microbial communities to specific nutrients. Prior studies have analyzed soil microbiome responses to essential nutrients that are assimilated into biomass, such as carbon, oxygen, nitrogen, and phosphorus, or biocides such as antibiotics and toxic metals (ref. 4, 5, 37, 85, 86). Our results show that a nanomolar amount of a biologically synthesized, non-assimilated nutrient can also cause rapid changes to the soil microbiome, underscoring the importance of corrinoids at the community level and the utility of our model nutrient approach. The requirement for multiple successive extractions to recover corrinoids from soil likely extends to other micronutrients, which are not typically observed in soil metabolomic analyses (ref. 63), limiting the complete characterization of soil metabolites. Taken together, the insights gained from this study form a roadmap for examining the functions of shared nutrients from individual microbes to complex communities, as well as to guide further study of corrinoid interactions in microbial communities in soil and other environments.

## Supporting information

Supplementary Table 1 - MAG Data

Supplementary Table 2 - Media Recipes

Supplementary Table 3 - Enrichment_16S_analysis

Supplementary Table 4 - Microcosm_16S_analysis

## Acknowledgements

This work was supported by the U.S. Department of Energy (DOE), Office of Science, Office of Biological and Environmental Research, Genomic Science Program under Award Number DE-SC0020155 to M.E.T., and National Institutes of Health award 5K99GM143653-02 to Z.F.H. Work conducted at Lawrence Livermore National Lab (LLNL), including soil microcosm incubations with H_218_O and the subsequent generation of genome-resolved metagenomes, was performed under the auspices of the DOE under Contract DE-AC52-07NA27344 with support provided by the DOE, Office of Biological and Environmental Research, Genomic Science Program LLNL ‘Microbes Persist’ Scientific Focus Area (award #SCW1632). We thank Katerina Estera-Molina for her expertise and management of the soil field site at Hopland Research and Extension Center. Peter Chuckran participated in helpful conversations about metagenome analyses. We thank Andy Goodman for providing the BtuG expression plasmid used in this work. We thank members of the Taga lab for helpful comments and critiques of this work. We thank L. Victoria Innocent for supplying [5-OHBza]Cba. We acknowledge that UC Hopland Research and Extension Center sits on the traditional, unceded land of the Pomo Indians. Research conducted at the University of California, Berkeley took place on the territory of xučyun (Huichin), the ancestral and unceded land of the Chochenyo-speaking Ohlone people, the successors of the sovereign Verona Band of Alameda County. This land was and continues to be of great importance to the Muwekma Ohlone Tribe and other familial descendants of the Verona Band.

## Data Availability

All 16S rRNA gene data generated or analyzed during this study are included in this published article [and Supp. Data Tables 3 and 4]. Metagenomic reads published previously in (ref. 37) and (ref. 38) are available in the NCBI SRA database under accession code PRJNA856348. Metagenomic reads and associated MAGs previously published in (ref. 36) are available in the NCBI SRA database under accession code PRJNA718849 and on ggKbase (https://ggkbase.berkeley.edu/wsip-metawrap-drep-bins/organisms).

