## Supplementary Figures for "Soil microbial community response to corrinoids is shaped by a natural reservoir of vitamin B_12_"

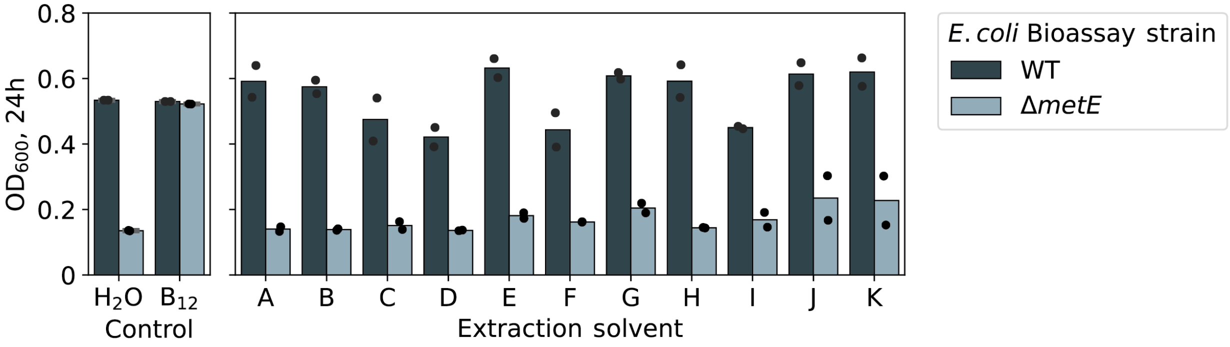


**Supplemental Figure 1. Quantification of total corrinoid in organic extractions from soil by microbiological assay.** Growth of the control *E. coli* MG1655 WT or corrinoid-responsive *ΔmetE* strains in M9-glycerol (0.4%) with added extract from soil obtained using the indicated organic solvent conditions. Solution A - MeOH:$H_2O$ (4:1), Solution B - $CHCl_3$:MeOH (2:1), Solution C - EtOAc, Solution D - Acidic MeOH:$H_2O$ (4:1), Solution E - Basic MeOH:$H_2O$ (4:1), Solution F - Acidic MeCN:MeOH:$H_2O$ (2:2:1), Solution G - Basic MeCN:MeOH:$H_2O$ (2:2:1), Solution H - MeCN:MeOH:$H_2O$ (2:2:1), Solution I - Acidic MeCN:$H_2O$ (4:1), Solution J - Basic MeCN:$H_2O$ (4:1), Solution K - MeCN:$H_2O$ (4:1). Acidic extraction solutions included 0.1 M formic acid, and basic solutions included 0.1 M ammonium hydroxide. CN-Cbl (vitamin $B_{12}$, 1 nM) or water ($H_2O$) were added as controls.

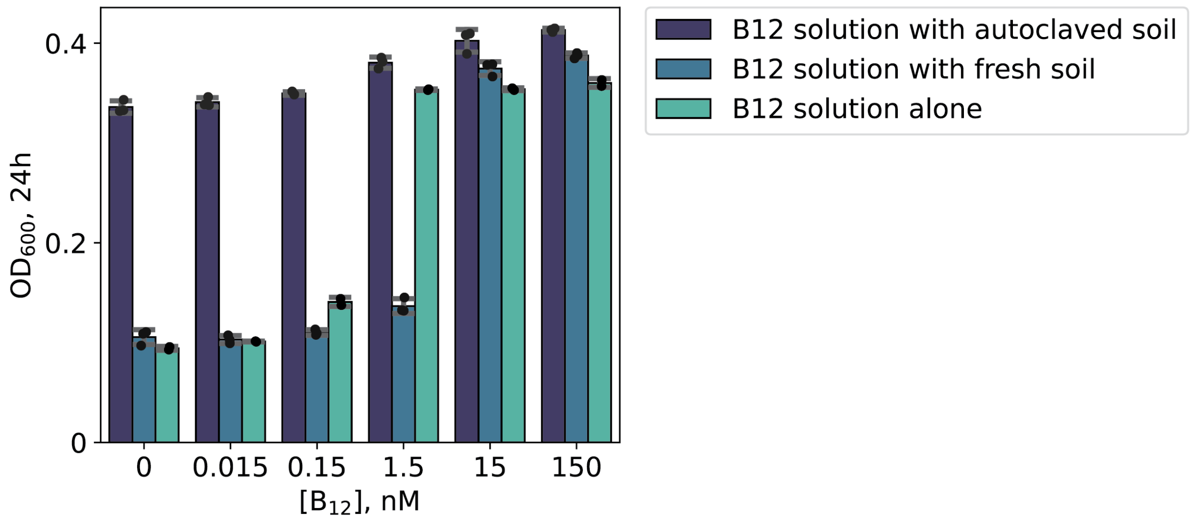


**Supplemental Figure 2. $B_{12}$ adsorbs to soil.** Growth of the corrinoid-responsive *E. coli ΔmetE* strain in M9-glycerol (0.4%) supplemented with CN-Cbl (vitamin $B_{12}$) solutions at the indicated concentrations pre-incubated with or without fresh or autoclaved soil.

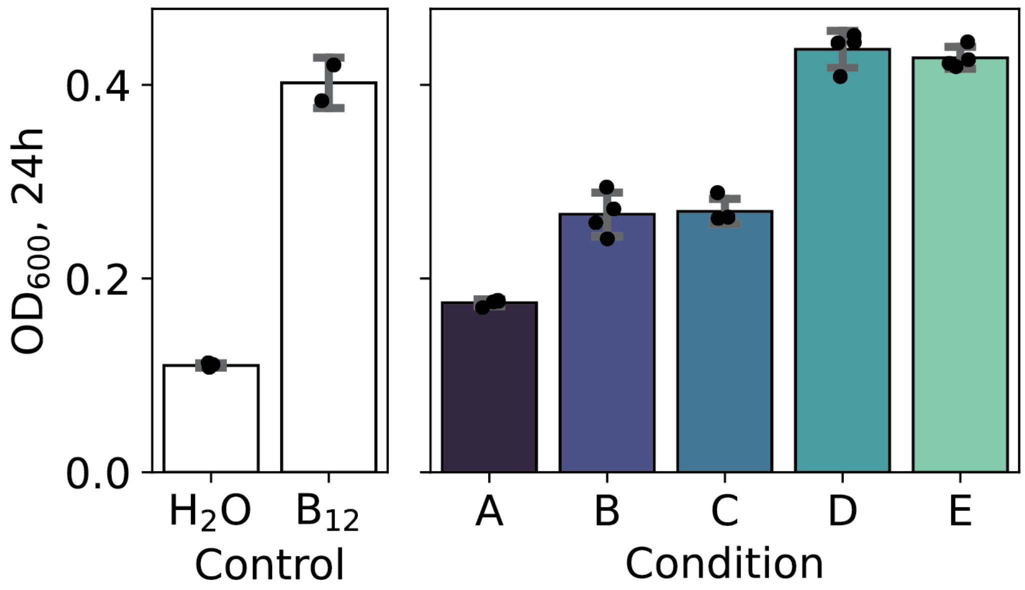


**Supplemental Figure 3. Effect of steam sterilization and phosphate buffer on corrinoid extraction from soil.** Growth of the corrinoid-responsive *E. coli ΔmetE* strain supplemented with control solutions of CN-Cbl ($B_{12}$, 1 nM) or water ($H_2O$), or extracts from soil: A - Fresh soil extracted with water, B - Fresh soil extracted with 0.1 M potassium phosphate (pH 7.0) solution, C - Autoclaved soil extracted with water, D - Autoclaved soil extracted with phosphate solution, E - Autoclaved soil that was moistened immediately before sterilization.

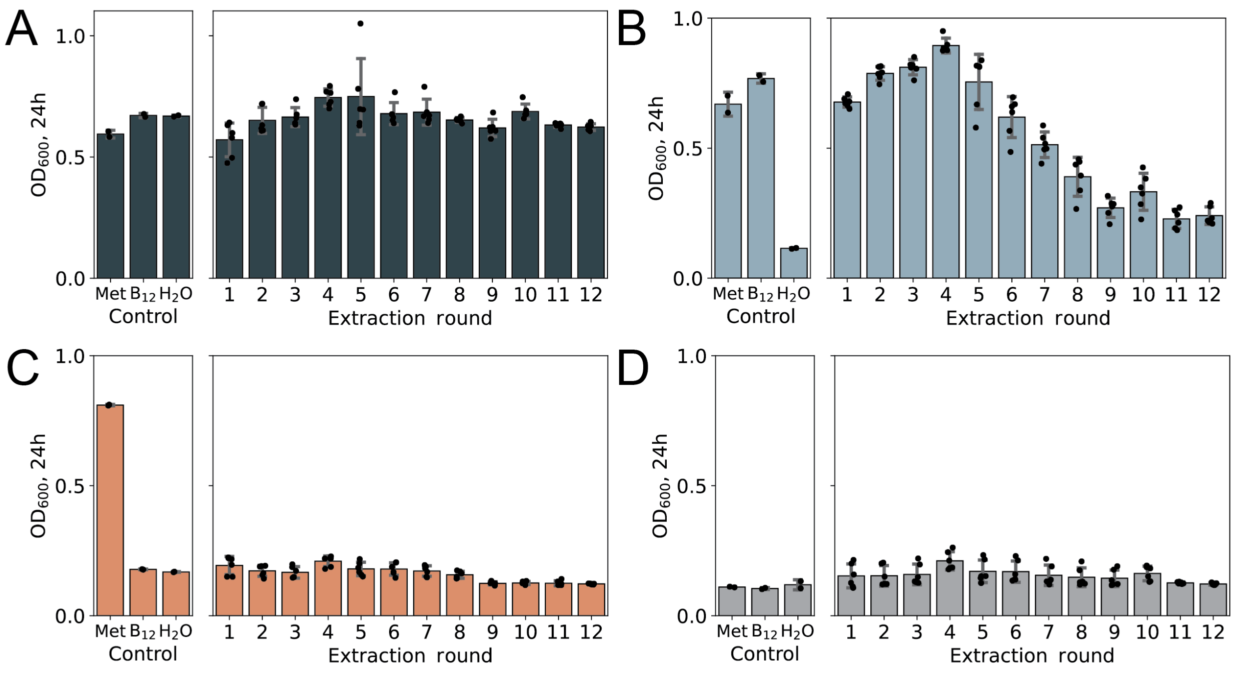


**Supplemental Figure 4. Controls for corrinoid quantification bioassay.** $OD_{600}$ measurements of *E. coli* strains: (A) WT, (B) *ΔmetE*, (C) *ΔmetEΔmetH*, and (D) uninoculated control in serial extraction rounds of soil using M9-glycerol (0.4%), alongside controls supplemented with methionine (Met, 6.7 mM), CN-Cbl ($B_{12}$, 1 nM), or water ($H_2O$). Panel B is a reproduction of Figure 3A, included here for readability.

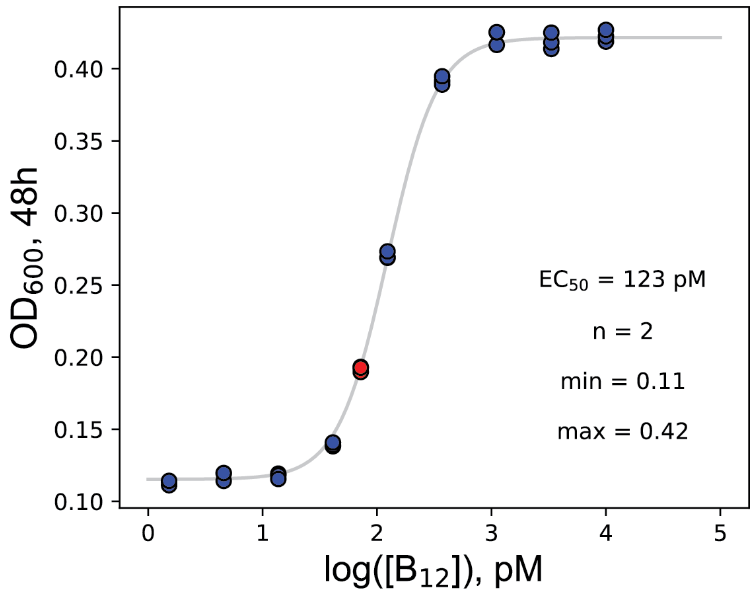


**Supplemental Figure 5.Quantification of total corrinoid in soil extracts in BtuG-depleted soil.** Dose-response curve of *E. coli ΔmetE* against CN-Cbl ($B_{12}$) in corrinoid-depleted soil extract (Blue), and corresponding measurement in control soil extract not depleted for cobamide by BtuG (Red). Soil extract used for this experiment was obtained by combining all 8 rounds of M9-based extractions (Figure 3A). Corrinoid-depleted soil extract was obtained by incubation with the high-affinity corrinoid-binding protein BtuG, followed by protein removal using Ni-NTA agarose beads. The control soil extract was treated only with Ni-NTA agarose beads.

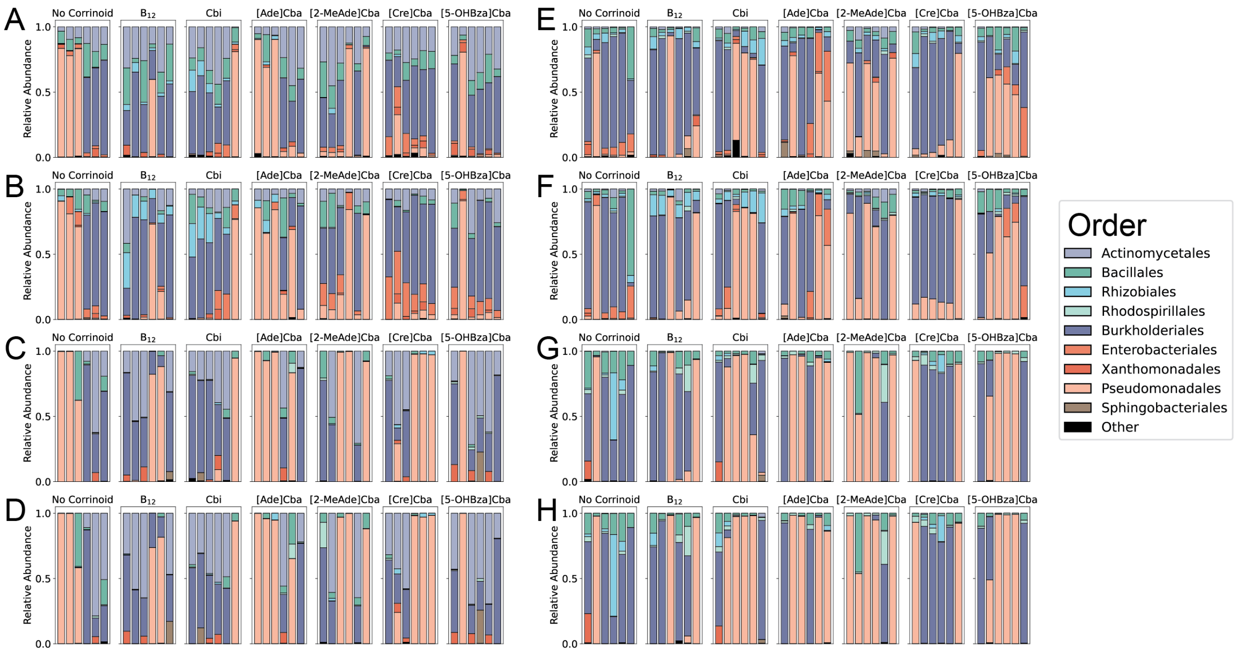


**Supplemental Figure 6. Relative abundances of taxa in soil-derived enrichment cultures based on 16S rRNA amplicon sequencing.** Relative abundances at the order level of enrichments in M9 (A-D) and R2 (E-H), grouped by corrinoid condition at (A,E) 1 week, (B,F) 2 weeks, (C,G) 12 weeks, and (D,H) 14 weeks. Six replicates are shown for each condition.

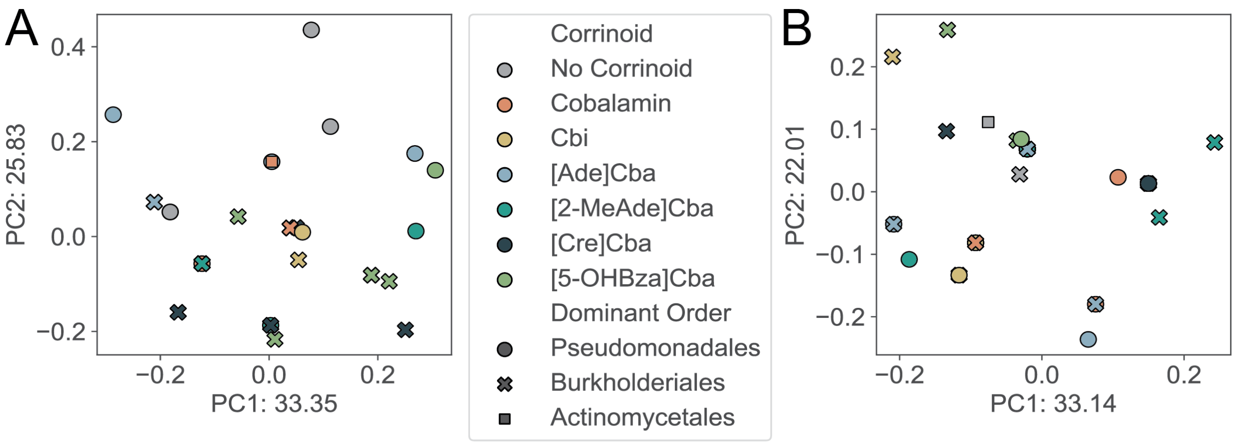


**Supplemental Figure 7. PCoA analysis using Jaccard dissimilarity metric of soil-derived enrichment cultures at 2 weeks.** A, B. Principal coordinate analysis (PCoA) plot with Jaccard dissimilarity of M9 (A) and R2 (B) enrichments at the order level grouped by corrinoid condition at 2 weeks. M9 enrichments (A) significantly cluster by corrinoid (ANOSIM: $r$ = 0.1189, $P$ < 0.05, PERMANOVA: $r$ = 2.407, $P$ < 0.05). R2 enrichment cultures (B) do not cluster by corrinoid (ANOSIM: $r$ = 0.3133, $P$ < 0.05).

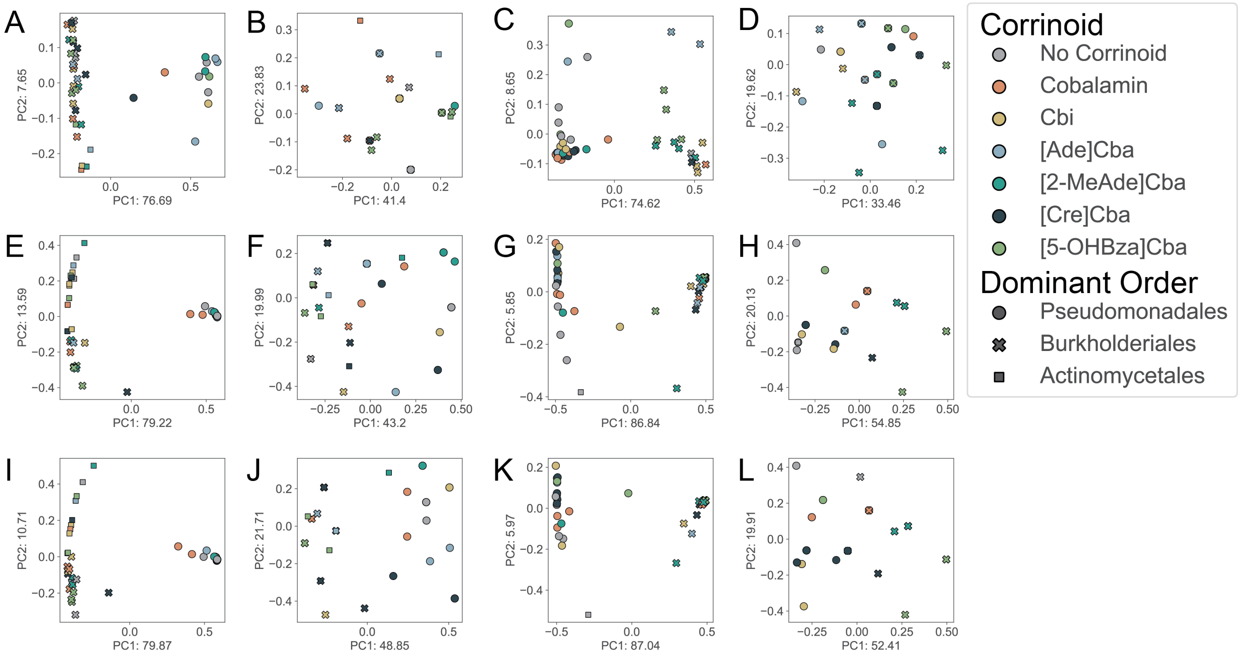


**Supplemental Figure 8. PCoA analysis of soil-derived enrichment cultures at 1, 12, and 14 weeks.** A-L. Principal coordinate analysis (PCoA) plot with Bray-Curtis (A,C,E,G,I,K) and Jaccard (B,D,F,H,J,L) dissimilarity of M9 (A,B,E,F,I,J) and R2 (C,D,G,H,K,L) enrichments at the order level at 1 (A-D), 12 (E-H), and 14 (I-L) weeks. No significant clustering is observed at these time points (ANOSIM/PERMANOVA, $P > 0.05$).

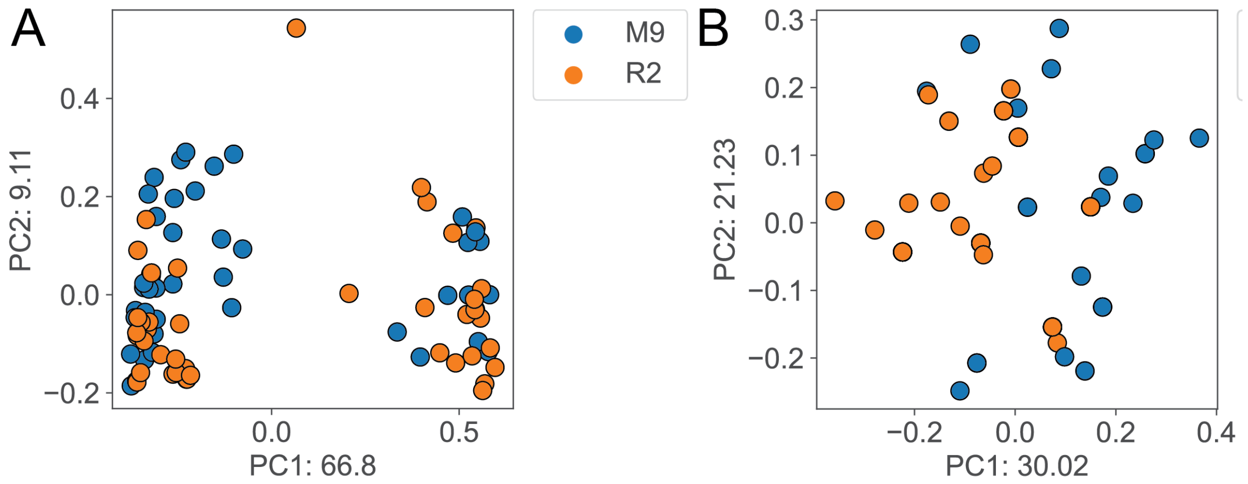


**Supplemental Figure 9. Clustering of soil-derived enrichments by culture medium.** A, B. Principal coordinate analysis (PCoA) plot with Bray-Curtis (A) and Jaccard (B) dissimilarity of M9 and R2 enrichments at the order level, grouped by medium (M9 and R2) at 2 weeks. Enrichments significantly cluster by medium using both (A) Bray-Curtis (ANOSIM: $r = 0.0652$, $P < 0.05$; PERMANOVA: $r = 3.8217$, $P < 0.05$) and (B) Jaccard (ANOSIM: $r = 0.3459$, $P < 0.05$; PERMANOVA: $r = 24.3267$, $P < 0.05$) metrics.

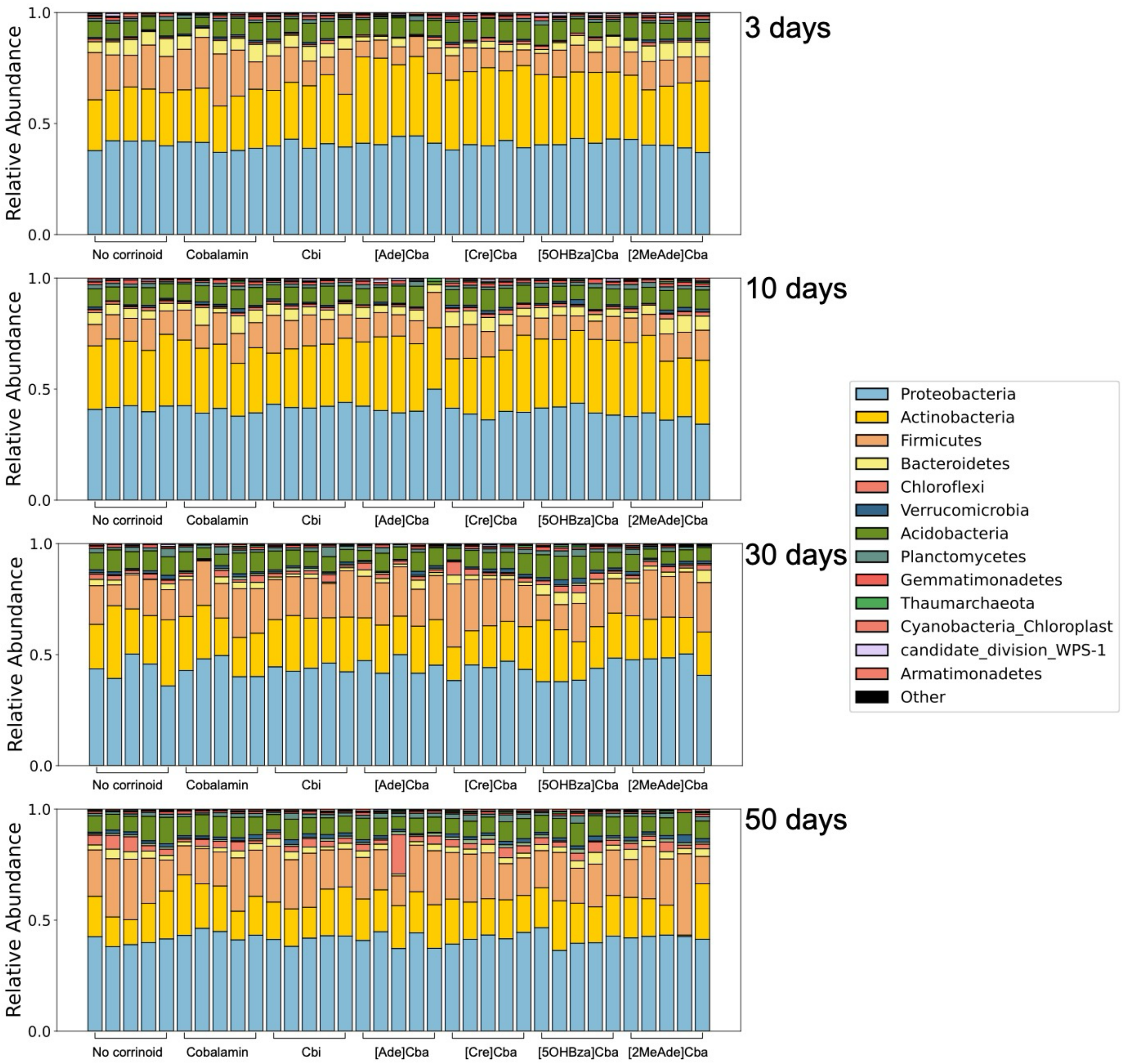


**Supplemental Figure 10. Composition of soil microcosms through time.** Stacked bar charts of the top ten phyla represented by colors, grouped by corrinoid treatment. Graphs are arranged through time where the top plot shows day 3 and the bottom plot shows day 50. Phyla are arranged with the most prevalent on the bottom and the least prevalent on the top.

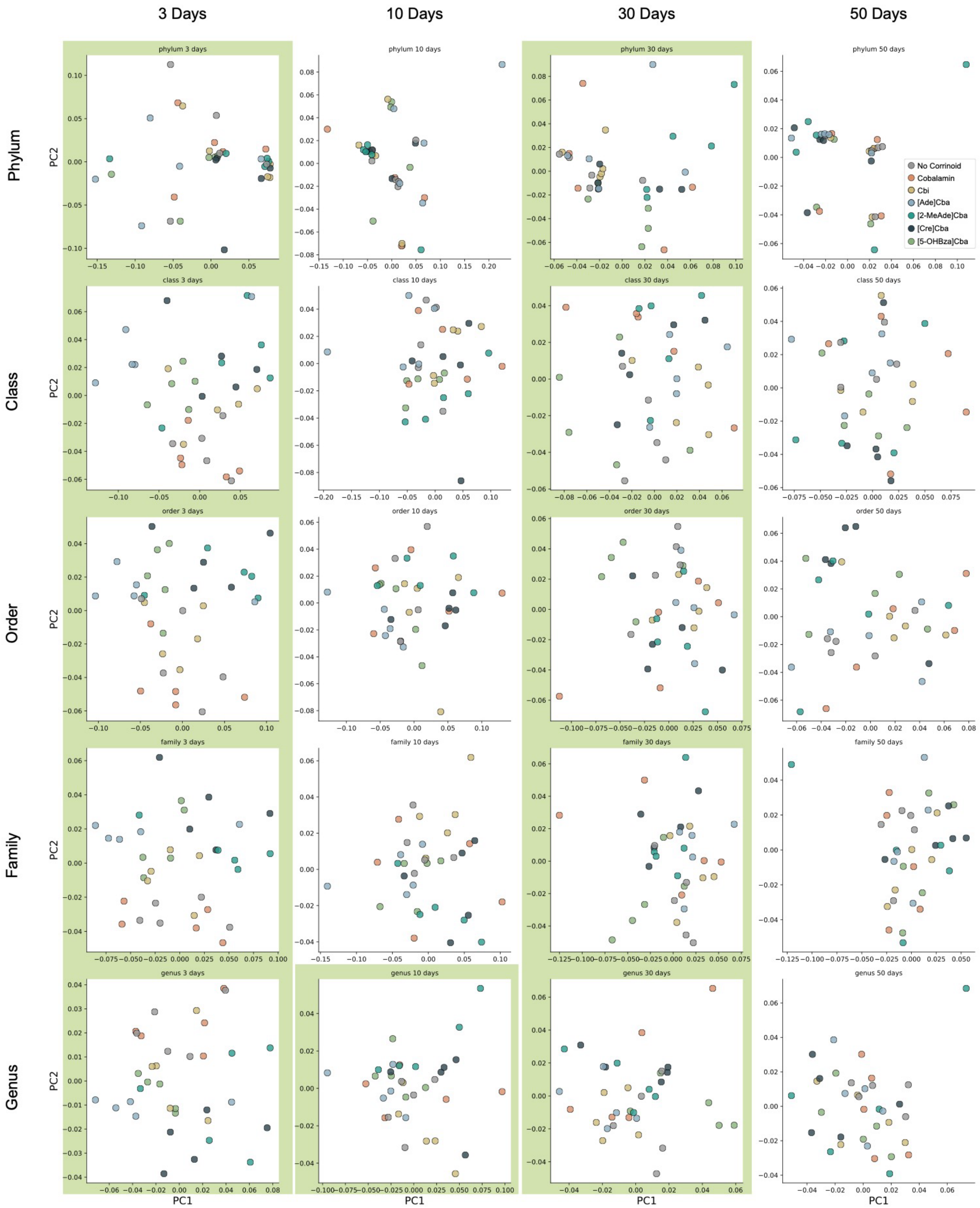


**Supplemental Figure 11. Corrinoid treatment effects on soil microcosms.** PCoA of Bray-Curtis dissimilarity aggregated by taxonomy (top: phylum, bottom: genus) through time (left: day 3, right: day 50). Each plot shows the first two principal components with the percent variance explained. Green shading indicates statistical significance (ANOSIM and PERMANOVA, $P \leq 0.05$).
